# Quantitative phase velocimetry measures bulk intracellular transport of cell mass during the cell cycle

**DOI:** 10.1101/2021.08.04.452408

**Authors:** Soorya Pradeep, Thomas A. Zangle

## Abstract

Transport of mass within cells helps maintain homeostasis and is disrupted by disease and stress. Here, we develop quantitative phase velocimetry (QPV) as a label-free approach to make the invisible flow of mass within cells visible and quantifiable. We benchmark our approach against alternative image registration methods, a theoretical error model, and synthetic data. Our method tracks not just individual labeled particles or molecules, but the entire flow of bulk material through the cell. This enables us to measure diffusivity within distinct cell compartments using a single approach, which we use here for direct comparison of nuclear and cytoplasmic diffusivity. As a label-free method, QPV can be used for long-term tracking to capture dynamics through the cell cycle.

## INTRODUCTION

Orderly transport is essential for cell function and growth. To maintain homeostasis, cells continuously transport materials, including nutrients and bulk material into cells from the surrounding environment^1^, ions through cell membranes^1^, and liquids^2^ and structural polymers into cell podia to drive cell motion^3^. In turn, cell transport can be impacted by disease^4^ and stress^5,6^. Measurement of transport within cells can therefore improve our understanding of cell behavior, disease, responses to environmental stresses and potential disease therapies. Fluorescent markers are the most commonly used tools to study transport within cells as they provide distinct signals with low background, making them easy to track^7-11^. Despite the wide use of fluorescent markers in transport studies, fluorescence come with the disadvantages of photobleaching and phototoxicity, which induce stress and modifies cell behavior^12,13^. The number of components labeled at a time is also limited^14^ and fluorescence tracking cannot measure velocities in untagged regions, limiting its ability to measure bulk material transport within cells.

Label-free methods offer an alternative to fluorescence for transport measurement. Label free methods, like phase contrast, differential interference contrast (DIC) and Raman imaging, have been applied to measure the dynamics of whole cells^15,16^, vesicles^17,18^, and lipid droplets^19^ within cells. However, DIC measures phase gradients in just a single orientation, and phase-contrast images contain halos that make quantitative analysis difficult^20^.

Quantitative phase imaging (QPI) is a label free imaging technique which measures the phase shift that occurs as light passes through a material with higher refractive index^21^. The phase shift measured with QPI is directly proportional to the distribution of dry mass in the sample^22^. QPI is, therefore, a better candidate for long-term transport studies within cells by providing quantitative data of the motion of bulk material within the cell. Previous work with QPI has used the contrast produced by localized variation in biomass density to track well-defined sub-cellular components as well as the overall average rate of mass motion^23,24^. Spatial and temporal power spectra from QPI data have also been applied to quantify cell-average rheological properties^25,26^, dynamics of red blood cells^27^, and localized diffusivities^28^.

In this work we combine automated image velocimetry and label-free QPI to develop an approach we call quantitative phase velocimetry (QPV) that measures unsteady intracellular velocity fields capturing the bulk transport of cellular material over long times. To understand the sources of error in QPV, we developed a theoretical model of QPV error that matches experimental results. We then apply QPV to measure dry mass transport inside cells during cell cycle progression. With this data, we quantify intracellular diffusion dynamics in the nucleus and cytoplasm over the cell cycle. We see nuclear diffusion decreases with cell cycle progression and cytoplasm diffusion reduces with G1 to S phase transition.

## RESULTS

### 1. Development of QPV from QPI data

We developed QPV to measure intracellular dry mass velocity from QPI data (**figure 1**). We capture QPI data using an inverted microscope with a quadri-wave lateral shearing interferometry (QWLSI) wavefront sensing camera^29^ (**figure 1a**) to obtain images of cell mass distributions over time. QWLSI uses a diffraction grating to cause interference between light that passed through adjacent regions of the sample, projecting an interference pattern onto the camera sensor. Wavefront aberrations, caused by the varying refractive index and thickness of the sample, are then captured as deformations of this interference pattern, from which the phase shift caused by the sample can be reconstructed^30^. QPI of retinal pigment epithelial (RPE) cells (**figure 1b**) shows the changing distribution of cell dry mass over time. In particular, QPI data show large regions of high mass density in nucleus, low mass podia, and small-scale puncta within cells that can serve as features for velocimetry (**figure 1b**). Taking the difference between QPI images over time illustrates the movement of mass captured by QPI (**figure 1c)**. In this example, this difference image indicates an overall movement of the cell dry mass from the top right corner to the bottom left corner of the image frame. QPV uses these data to measure the movement of mass using the principles of particle image velocimetry (PIV). As expected in this individual, example cell, the distribution of intracellular dry mass velocity determined by QPV shows dry mass velocity vectors pointing from the top right to the left bottom of the frame, as well as localized deviation from that overall trend (**figure 1d**).

**Figure 1.**
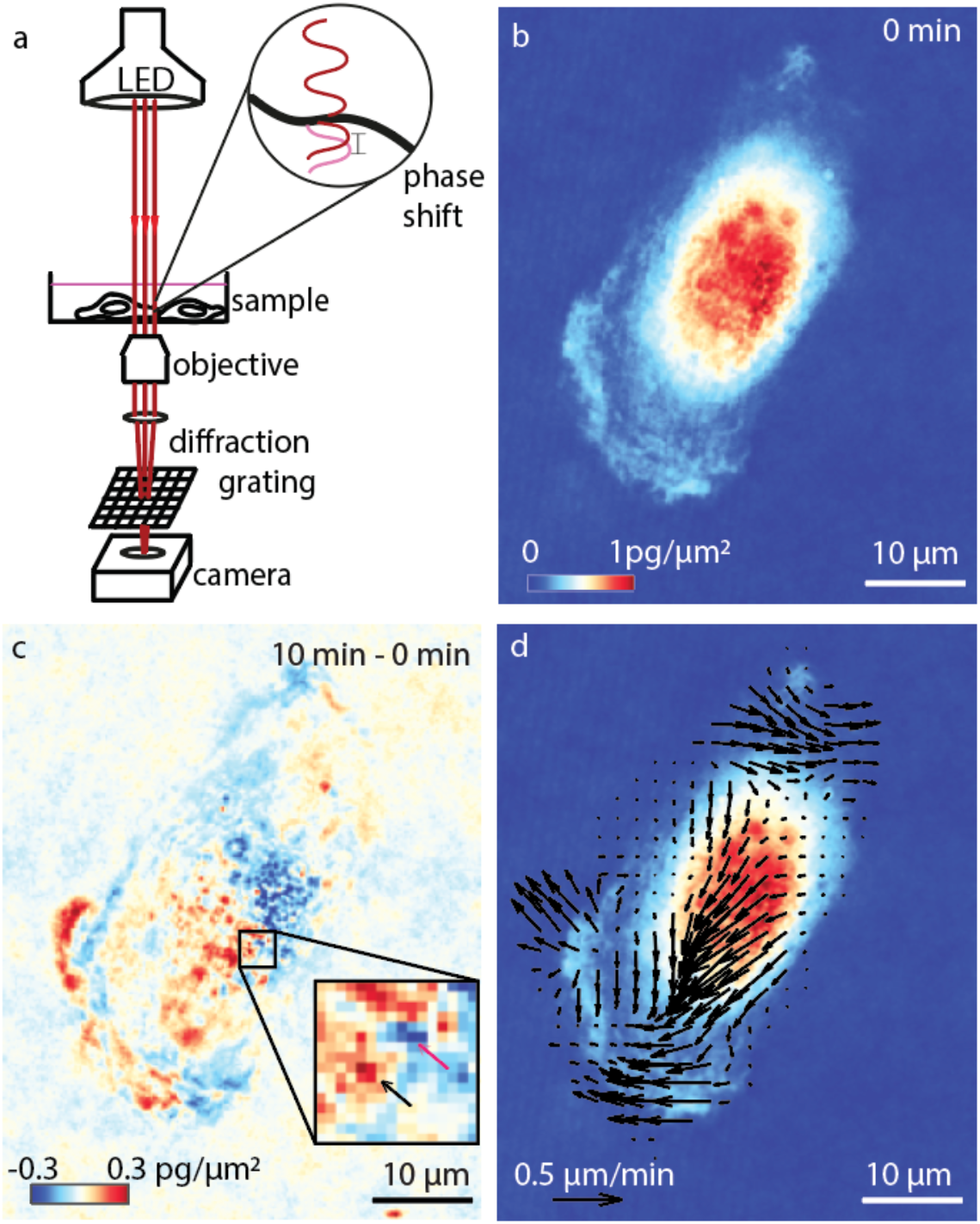
Quantitative phase velocimetry (QPV) measures intracellular dry mass movement. (a) Quantitative phase imaging (QPI) measures the phase shift of light passing through a cell, which is used to compute the dry mass distribution in cells over time. (b) Dry mass distribution in RPE cell imaged at 120X magnification, at *t* = 0 min. The scalebar indicates 10 µm length. (c) The difference in QPI mass distribution of the RPE cell in (b) from an image taken at *t* = 10 min later minus the image at 0 min reveals cell motion. The color scale shows the net displaced mass over this interval (red increase, blue decrease). The inset in (c) shows a 15×15 pixel interrogation window that illustrates the change in position of an individual subcellular feature from the position marked with a red arrow to the position marked with a black arrow. Colorbar shows dry mass difference between frame at time 0 and 10 minutes (red: large increase in mass, blue: large decrease in mass). (d) The resulting intracellular biomass velocity field computed using quantitative phase velocimetry (QPV). Velocity magnitude indicated with a 0.5 µm/min scalebar.

To choose the most compatible image registration method for developing QPV we compared the performance of methods commonly used for PIV: normalized cross-correlation (NCC)^31^, optical flow reconstruction (OFR)^32^, mutual information (MI)^33^, and sum of squared differences (SSD, **figure S1**)^34^. With each method, we estimated the resolution and accuracy of velocity measurement on fixed RPE and MCF7 cells moving at known velocity as a velocity standard, as fixed cells have the same distribution of intracellular features as living cells (**figure 2** and **figure S2-S3**). The displacement distribution computed by QPV of example fixed cells after a total of 1.5 µm downward stage motion shows uniform displacement distribution in all interrogation windows pointing in the direction of stage motion (**figure 2a** and **figure S2**).

**Figure 2.**
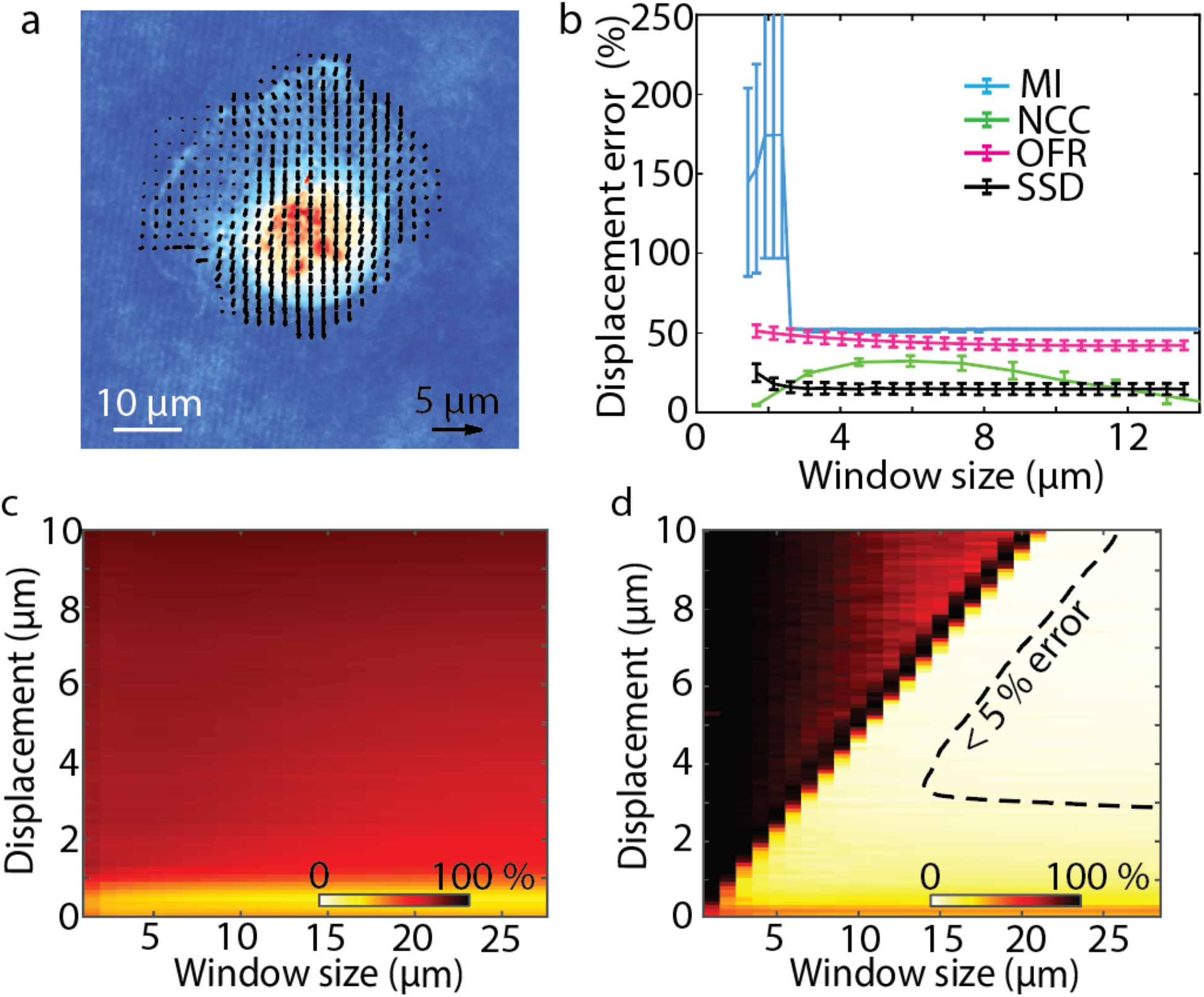
SSD image registration measures intracellular velocity from QPI data with higher accuracy than OFR, MI and NCC at most interrogation window sizes. (a) QPV on RPE fixed cell measures the intracellular displacement during microscope stage translation. (b) Comparison of displacement error, the percentage difference between the measured and expected displacements, for mutual information (MI), normalized cross correlation (NCC), optical flow reconstruction (OFR) and sum of squared difference (SSD) for intracellular velocity computation on QPI data at different interrogation window sizes shows SSD has highest accuracy at most window sizes (number of cells, *n*, = 11, error bar shows standard error of the mean, SEM). (c) Velocity accuracy versus both interrogation window size and displacements of RPE fixed cell using OFR shows that OFR is limited to measuring a very narrow range of displacements with acceptable accuracy (*n* = 11). Yellow/white – high accuracy, red/black – low accuracy. (d) QPV velocity measurement accuracy versus interrogation window size and displacements of fixed RPE cells show typically less than 10% error when the interrogation window size is smaller than the displacement to be measured with a region of less than 5% error indicated as a dashed line (*n* = 11).

We need a small interrogation window size for our velocimetry approach to achieve high spatial resolution of the velocity field. We computed the measurement accuracy of 0.1 µm (0.42 x a single pixel) displacements of fixed cells using each method with interrogation window sizes ranging from 5 to 59 pixels (1.19 µm to 14 µm) square. We computed displacement error as the percentage difference of the measured displacement from the expected displacement (see Methods). The displacement error is the key determinant of the error in computed velocity as the time between frames is tightly controlled during imaging. Of the four methods tested, SSD, NCC and OFR worked fairly well for all window sizes, and MI worked moderately well with large windows (**figure 2b** and **figure S3a)**. The similar performance of SSD and NCC is expected, as the SSD is related to the cross correlation (see Methods). However, the cross correlation alone shows inferior performance to either SSD or NCC due to the nonuniform, background energy density of QPI data (**figure S4**). Though MI and OFR have large errors in application to QPI data, they have reduced computation time relative to NCC and SSD. SSD in particular shows a reduction in computational time with smaller interrogation window size (**figure S5**).

The image registration method used for QPV should also perform well over a wide range of displacements as cells typically display a wide range of motion from sub-pixel movement in the nucleus to highly deforming multiple pixel displacements in the podia regions of the cytoplasm. Based on their performance at small to medium window sizes and basis in fundamentally different image registration methods, we further measured displacement error with SSD and OFR for displacements from 0.1 to 10 µm and interrogation window sizes from 1 to 14 µm. The results for OFR with fixed RPE and MCF7 cells shows greater than 50% error for displacement measurements above 1 µm at all interrogation window sizes tested (**figure 2c** and **figure S3c**). Performance of OFR at large displacements can be improved using Gaussian blurring^35^ (**figure S6**). However, Gaussian blurring comes at the cost of losing the ability to resolve small-scale differences in deformation within cells. On the other hand, SSD gives an error of less than 10% for any displacement, as long as the window size for the calculation is at least as large as the displacement itself and less than 5% for a subset of this region (**figure 2d**). We also evaluated SSD has less error than OFR in displacement direction measurement (**figure S7**). SSD also performs better than OFR for different cells (**figure S8**). Therefore, considering accuracy, spatial resolution and the ability to measure a wide range of displacements, and despite a moderate increase in computational time (**figure S5**) we chose SSD as the image registration method for QPV.

### 2. Theoretical modelling of displacement measurement accuracy

To understand the sources of errors in QPV, we developed a model that account for key parameters that contribute to measurement noise. These are: the size range of cell features which impacts Brownian diffusion of cell components and the ability to visualize displacement with the interrogation window size chosen, the interrogation window size which determines the maximum measurable displacement, and the optical resolution limit of the microscope.

To study how the distribution of spatial features and noise influence the performance of QPV, we generated synthetic data consisting of uniform circles (**figure 3a** top middle) and non-uniform-sized circles (**figure 3a** top right) with sizes matched to the average particle sizes observed in RPE and MCF7 cells (**figure 3b** and **figure S9i)**. The Perlin noise added to synthetic data matches the continuous local variation background noise in our QPI data (**figure 3a** bottom middle and bottom right). Perlin noise thus has similar power spectrum as the background noise in QPI, and reduces the magnitude of the power spectral density (**figure S9i**). The synthetic Perlin background noise gives results comparable to addition of noise from real QPI images (**figure S9i**). The use of synthetic Perlin noise then creates for a completely synthetic dataset that independent of the real images was used to both determine the essential characteristics that limit the measurement and to make an easily generated synthetic dataset for evaluation of the methods. The power spectra of the RPE fixed cell, MCF7 fixed cell, and polystyrene beads imaged at 20X magnification match best with the power spectrum of non-uniform synthetic data with Perlin noise (**figure 3b** and **figure S9i**). Therefore, based on this power spectrum analysis, QPI images are best approximated as having as non-uniform spatial features with added low spatial frequency noise. QPV on synthetic data shows less than 5% error for both data with uniform and non-uniform spatial features (**figure S10**).

**Figure 3.**
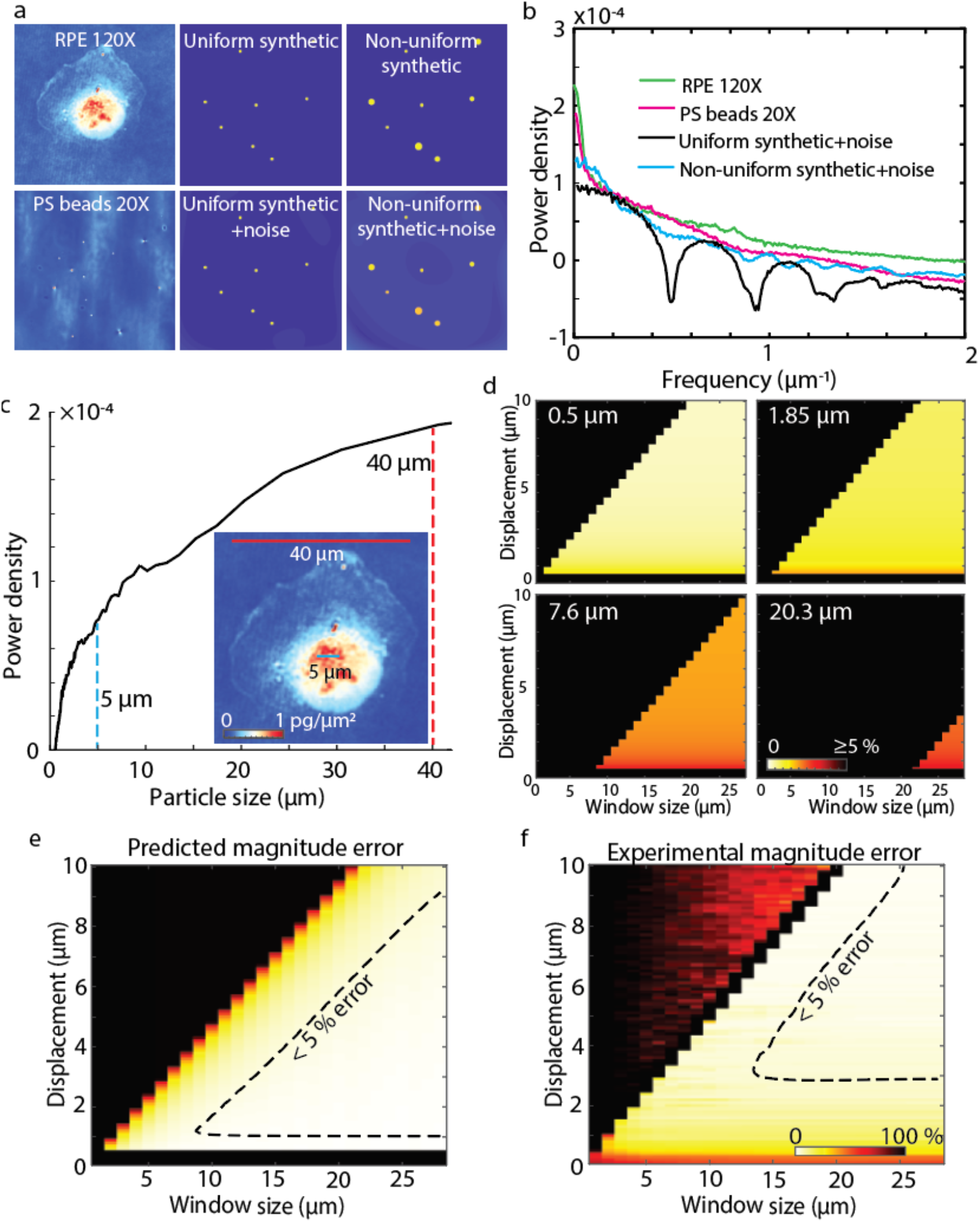
Modelled QPV displacement measurement error agrees with experimental measurements. (a) Comparison of background corrected QPI image of RPE fixed cell at 120 X (top left) and polystyrene beads at 20X (bottom left) to synthetically generated uniform and non-uniform circle data, with added Perlin noise (top middle and right) and without added Perlin noise (bottom middle and right) (b) Power spectrum of the QPI data (green and magenta) and synthetic data with noise (black) in (a) shows the similarity of QPI data to the non-uniform circle synthetic data. (c) Power spectral density versus effective particle size of RPE fixed cells (*n* = 3). Power spectral density corresponding to both 5 µm and 40 µm structures is indicated. Sub-panel shows structure inside 40 µm RPE fixed cell illustrating the range of features captured by QPI. Scale bars show 5 µm (blue) and 40 µm (red). Colormap shows dry mass density in pg/µm^2^. (d) Theoretical displacement estimation error (percentage difference between expected and measured displacements) at four effective particle sizes for all tested displacements and window sizes. (e) Averaging the size dependent model results as in (d), weighted by the actual distribution of particle sizes (c) gives a predicted error which is in agreement with the (f) measured error from the matched RPE cell using experimental data. Colormaps in (e) and (f) are the same.

Converting frequency to equivalent particle size shows that the features potentially able to be tracked by QPV range in size from organelle-size puncta of mass to the entire cell (**figure 3c**). To account for the range of feature sizes captured by QPI and their individual contributions to measurement error, we compute the theoretical error for all particle sizes in a cell up to 40 µm, the size of a whole cell (**figure 3d, figure S11c-d**), then perform a weighted average of these particle errors based on the power spectrum for the cell (**figure 3c, figure S11b**). The resulting prediction of the error in displacement magnitude (**figure 3f, figure S11e**) are in good agreement with the experimentally computed error for both RPE (**figure 3e)** and MCF7 cells (**figure S11f**). Both predicted and measured error show large error if the displacement is larger than the window size as well as large error when the displacement is less than the diffraction limit (0.48 µm in our case).

We achieve diffraction limited optical resolution by adjusting the phase pixel size to be half that of the diffraction limit, therefore satisfying the Nyquist criteria (**figure S12a-c**). We then observe high velocity estimation error at displacements below the optical resolution limit of the microscope in the experimental (**figure 3f**) and predicted (**figure 3e**) displacement magnitude. To estimate the impact of optical sectioning on our velocity estimation, we first compute the depth of field of our system as 1.7 µm, which is fairly large due to the use of low condenser aperture and large effective pixel size. We can then use the calculate the optical thickness of the sample, called optical path length (*OPL*), to estimate the thickness of the cells we imaged with QPV. Assuming a refractive index of DMEM cell culture media as 1.337^4^ and a typical refractive index for cell contents of 1.36^5^, we estimate that cells with OPL less than approximately 0.04 µm will be within the system SOF. We observe that the maximum optical path length in RPE cells is mostly within this range (**figure S12d**). Additionally, we note that, while some features which are beyond the depth of field will be observed as blurred features, previous work on PIV indicates that this will not have a large impact on velocity estimation accuracy^36^. We also observe reasonable agreement in the predicted and measured displacement direction error (**figure S13**). Application of this model and validation allows us to predict the minimum window size for accurate intracellular velocimetry. For example, for RPE and MCF7 cells imaged at 120X magnification every 1 min, the typical observed range of displacement varies from 0 to 14 pixels outside of the very fast moving podia. Therefore, we use a 15 by 15 pixel interrogation window, which corresponds to an area of 12.75 µm^2^. We note that QPV can be adapted to even faster moving cells by increasing the interrogation window size or by increasing the rate of imaging, therefore capturing larger movements between shorter time intervals with reasonable error down to motions that approach the diffraction limit when suitable optics are used.

QPV accuracy also depends on the cell type and varies based on feature density, which indicates the effective particle count tracked by QPV and varies across different regions within cells (**figure S12**). The error in intracellular velocity in an example RPE and (**figure S14a-c**) MCF7 fixed cell (**figure S14d-f**) are higher towards the edge of the cells with flat featureless regions. Additionally, the lower power spectral density in RPE cells relative to MCF7 cells (**figure S14g**) indicates that the power density of larger particles is more dominant in RPE cells. This which moderately reduces the accuracy of QPV. The velocity at different interrogation windows in cells also shows increased error in windows with lower power density (**figure S14h-i**).

### 3. Application of QPV for measurement of intracellular displacements

QPV tracks both overall cellular deformation as well as the motion of individual regions within cells. As a method based on QPI, which measures dry mass distributions^22^, QPV measures displacement of dry mass. Sample results for an RPE cell are shown in **figure 4**. The deformation of a uniform grid overlayed on the RPE cell image from time 0 minutes (**figure 4a**) to 30 minutes (**figure 4b**) shows compression of the grid points in the nuclear region and the spacing out of points in the podia region (**figure 4b**), reflecting the observed compression of the cell as its podia move upward within the image frame (**movie S1**). Each square enclosed by four grid points in **figure 4a** represents a volume encompassing a specific quantity of dry mass in the cell. Using QPV, we track each region of dry mass as it moves and deform with time. QPV, therefore, quantifies the intracellular velocity magnitude and direction in each volume inside the cells, including in high velocity podia regions (**figure S15**). The motion of five such volumes in the nuclear and cytoplasmic regions of the indicated RPE cell shows upward movement of the control volumes and the movement of the cell over 30 minutes (**figure 4c-d** and **movie S2**). From these tracks, we can see that mass originating from the central region of the cell are more direction oriented with smaller scale fluctuations than mass originating in the lower density regions of the cytoplasm.

**Figure 4.**
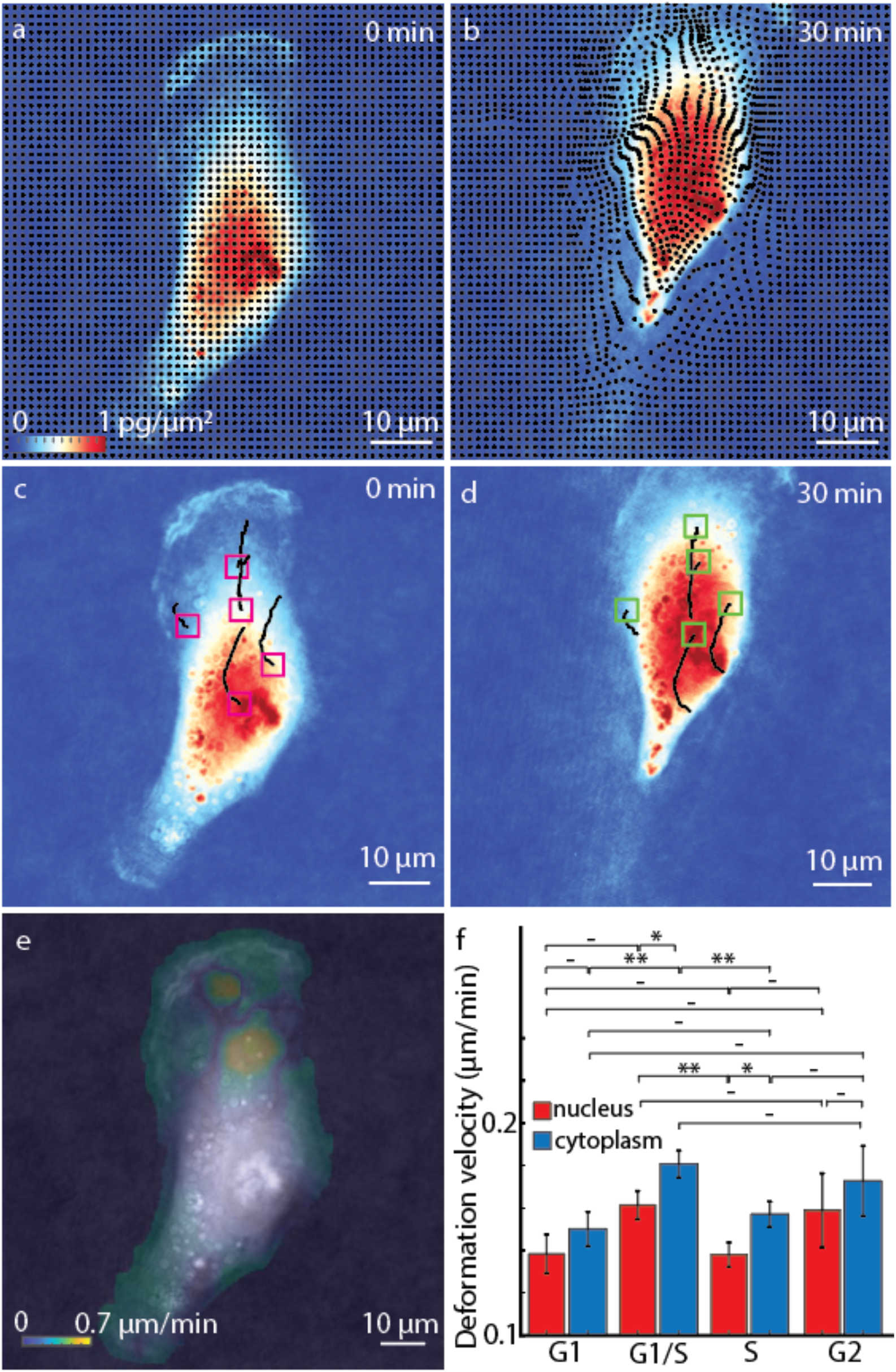
QPV shows spatial and temporal dynamics of biomass motion within cells. (a) Grid markers depict 4 by 4 pixel intracellular volume centroids overlayed on an RPE cell at 120 X magnification (b) which deforms from (a) to (b) due to movement of the cells in 30 minutes. (c) Dry mass inside control volumes initial positions marked using magenta boxes travel along the black line to reach the final positions indicated using green boxes in (d) in 30 minutes. (e) Deformation velocity, the whole cell velocity subtracted intracellular velocity distribution of dry mass, inside the RPE cell, gives the magnitude of the deviation of each cell volume’s motion from its expected position. (f) Deformation velocity of dry mass in nucleus and cytoplasm of RPE cells, indicating magnitude of the local cell deformation, shows higher deformation in cytoplasm than nucleus (*n* = 59, error bars show standard error of the mean). *\* p < 0*.*05*, ** *p* < 0.01.

QPV data quantifies the overall velocity of bulk mass transport of each region within the cell. The mean velocity distribution over 30 minutes in the cell shows a higher average velocity in the cytoplasm region, with a moderate velocity in the nuclear region (**figure S16a**). To more systematically assess differences between the cytoplasm and nucleus, we segmented the nuclear region from the cytoplasm of the RPE cell using the fluorescent images of an expressed FUCCI marker^37,38^. Separating the velocities in the nucleus and cytoplasm and comparing them to the cell centroid velocity showed no difference as the cell nucleus and bulk of the cytoplasm moves with the cell (*n* = 59 cells, **figure S16b**) and maximum velocities of up to ∼1 µm/min (**figure S16c**). Over the 1 min imaging interval, these maximum velocities correspond to displacements of up to 14 pixels, just within the acceptable error bounds determined by our error characterization (e.g. **figure 3**). We compute the deformation velocity as the velocity of the control volumes within the cell relative to the cell centroid velocity. The deformation velocity computes the magnitude of the deviation of each cell volume’s motion from its expected position if the cell moved as a rigid body. It is defined as the difference between each intracellular volume motion relative to the overall movement of the cell centroid (which is typically tracked, e.g. in motility studies). The deformation velocity map overlayed on the RPE cell (**figure 4e**), shows a similar distribution as the overall distribution of intracellular velocities **(figure S16a**). However, the velocity magnitude in the regions showing the smallest deformations, such as the nucleus, are reduced to around zero (**figure 4e**). The average deformation velocity from the nucleus and cytoplasm of the cell also reflects a larger deformation in the cytoplasm compared to the cell nucleus (**figure 4f**). Thus, the nuclei of RPE cells move more in line with the overall cell body relative to the cytoplasm, with the podia in particular exhibiting a larger velocity relative to that of the whole cell. We also observe higher activity in the cell nucleus and cytoplasm during G1 to S phase transition (**figure 4f**) perhaps due to the increased G1-S transcriptional activity to prepare for the DNA replication^39^.

### 4. Decreasing intracellular diffusion with cell cycle progression

Using QPV, we tracked the intracellular dynamics of mass within every control volume within the cell (**Figure 4c-d**) to measure transport of mass continuously for eight hours while simultaneously monitoring cell cycle progression with the FUCCI cell cycle indicator^37^. From these data, we then performed the mean squared displacement (MSD) analysis of the deformation, or displacement relative to cell centroid displacement, of all tracked intracellular mass from QPV of RPE cells in cell cycle (**figure 5**). The slope and intercept of the MSD plot was used to measure the anomalous constant and diffusion coefficient, respectively, of each intracellular region tracked with QPV (**figure 5a-c**). To validate our MSD calculation, we compare the anomalous and diffusion constants of live RPE cells to fixed cells moved artificially on a stage that show effectively zero diffusivity (**figure S17**). Nuclear boundaries were determined by alignment to fluorescence images of FUCCI cell cycle markers (**figure 5b, figure S18**). These data indicate that although cytoplasmic material exhibits a larger range of displacements than material within the nucleus (**figure 5e-f**), these displacements indicate a lower average effective diffusivity (**figure S19a**). Both nuclear and cytoplasmic diffusion were consistent with moderately sub-diffusive (**figure 5g**), anomalous diffusion (**figure 5h**).

**Figure 5.**
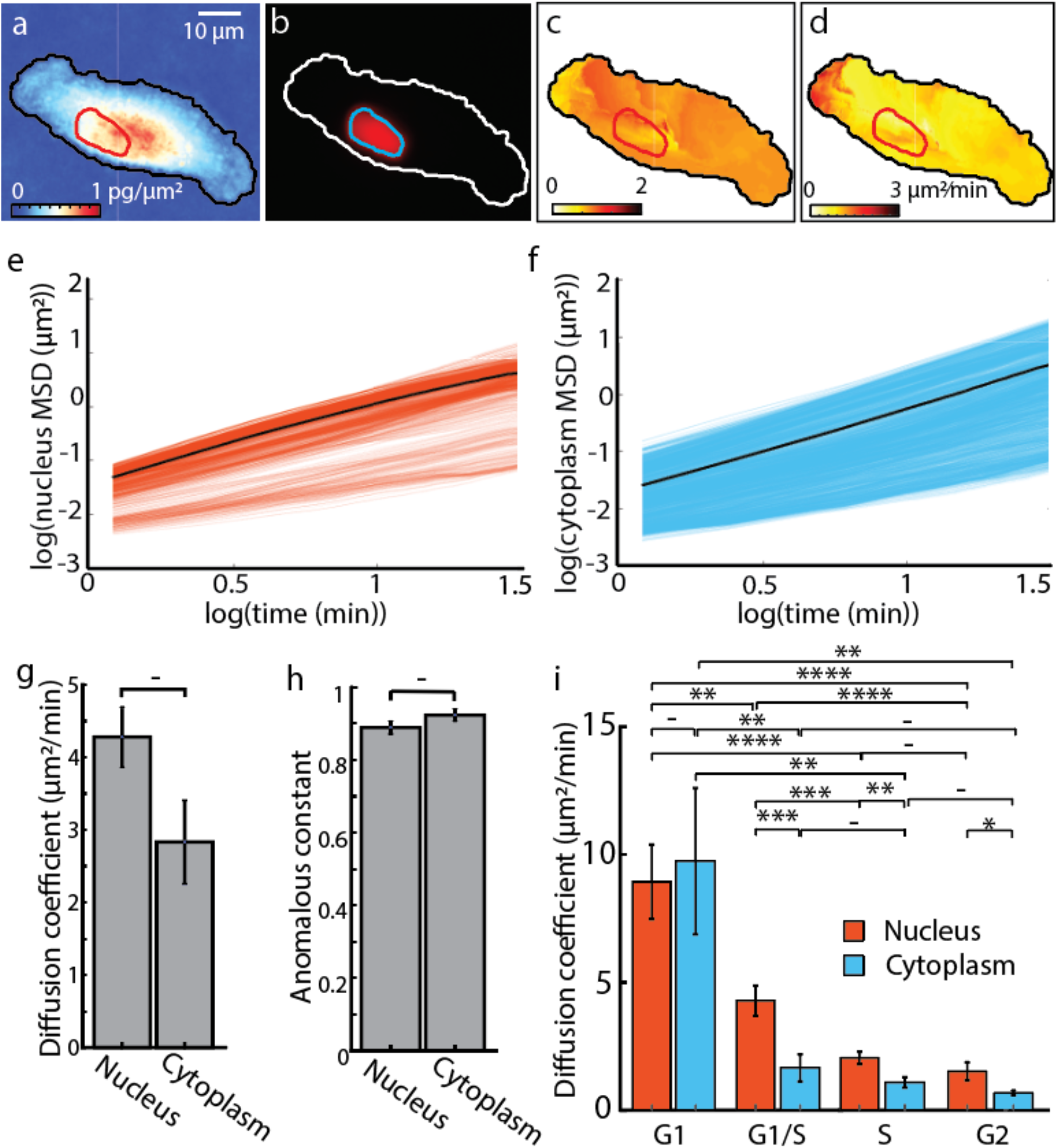
Mean squared displacement (MSD) analysis shows reduced diffusion coefficient in RPE cells with cell cycle progression. (a) QPI image of individual RPE cell with red nuclear and black cytoplasmic boundary shown. (b) Anomalous constant distribution within the RPE cell from (a). (c) Diffusion coefficient distribution within the RPE cell from (a). (e) Logarithm of MSD vs time lag of nuclear volumes of the RPE cell in (a). The black line shows the mean of all nuclear volumes. (f) Logarithm of MSD vs time lag of intracellular cytoplasm volumes of RPE cells in (a). Black line shows the mean of all cytoplasmic volumes. (g) Separation of diffusion coefficient in nucleus and cytoplasm of the RPE live cells shows moderate, but not significant reduction in diffusion coefficient in the cytoplasm relative to the nucleus. (h) The anomalous constants from the MSD analysis on the same cells shows moderately sub-diffusive transport in both the nucleus and cytoplasm. (i) Nuclear and cytoplasmic diffusion coefficients in RPE cells over the cell cycle from *n* = 119 cells. Error bars show standard error of the mean. – *p* > 0.05 (not significant), ** *p* < 0.01, *** *p* < 1×10^−3^, **** *p* < 1×10^−4^.

The average diffusion coefficient was slightly higher in the nucleus than in the cytoplasm with wide variation (**figure 5g**). To understand this variation, we binned the diffusion coefficient based on cell cycle phase (**figure 5g, figure S19**). These data indicate that the nuclear diffusion coefficient reduces with cell cycle progression and cytoplasm diffusion coefficient reduces through G1 to the beginning of S phase and agree with estimates of spatially variable intracellular diffusivity over similar time scales using quantum dots^40^. The range of diffusion coefficient also is similar to the range estimated using genetically encoded multimetric nanoparticles (GEMs)^41^, nanosized particles that freely move through the cytosol and nucleus. However, QPV but shows the opposite trend of cytosol diffusion coefficient being higher than nucleus. This may be expected as QPV includes the diffusion of organelle networks inside cells and not the liquid flow, thus reducing the effective diffusion coefficient values. As a method that tracks mass of intracellular components, we can apply QPI to determine the dependence of the measured diffusivity of each control volume on its mass (**figure S20a**), and separately perform the same analysis in different compartments of cells such as the nucleus (**figure S20c**) and cytoplasm (**figure S20b**). This is close to the expected scaling of D ∼ m^-1/3^ that would be predicted from Stokes-Einstein, assuming mass scales with the cube of effective particle size, but with significant deviation at the level of individual control volumes. We estimated the effective size of particles tracked by QPV from the power spectra of QPI images (**figure 3a**), limited to particles that are within the window size used for QPV. This yields a roughly constant effective particle size through the cell cycle (**figure S19b**).

## DISCUSSION

We developed QPV to measure unsteady velocity fields of bulk intracellular transport from QPI. In contrast to bulk transport across the cell membrane and into cells, the bulk intracellular transport measured with QPV is the transport of all cellular material within the cell. As a label-free method, QPV can directly be applied to measure mass transport within cells over long periods, such as throughout the cell cycle, and to make direct comparisons between cellular compartments. We also applied QPV to quantify the transport properties of subcellular trajectories to measure intracellular effective diffusivity through the cell cycle.

Our model of error contributions to QPV points towards possible improvements. One key insight is that the particle size distribution is critical in determining the accuracy of QPV. This suggests that a high-pass filtering scheme may improve accuracy of QPV by reducing the effective particle size. However, this would come at the loss of applicability to tracking whole cell displacements, as the resulting velocity fields would be for a subset of intracellular particles. We also note that, though Brownian motion was found to be a key limit to the accuracy of velocity estimation for previous applications of PIV to microscopy data to measure flow within microchannels^42,43^, we found the influence of Brownian motion to have a small impact on QPV. We estimate an impact of <3% for the smallest particle sizes imaged and negligible impact for the largest particles tracked. We also note that as a method based on image registration rather than particle tracking, QPV should be able to handle some degree of particle disappearance and reemergence e.g. due to fusion and fission. However, in regions of low feature density, where QPV error in fixed cells was found to typically be higher than in feature-dense regions, the appearance/disappearance of individual particles will add further error in living cells.

While our demonstration focused on the QWLSI method for QPI, there are a wide range of alternative methods available for obtaining 2D or 3D QPI data that could potentially be analyzed with QPV. For example, other 2D QPI methods such as digital holographic microscopy ^44^ that capture a single phase pixel per camera sensor pixel could potentially increase the number of pixels available for analysis. This would increase the possible interrogation area (e.g. allowing imaging of more than one cell, or cells with larger spread area) and increase computational cost accordingly. QPV can also potentially be extended to use with 3D QPI data available from 3D tomographic QPI imaging^45,46^. This would come at the cost of increased computation time. As we have found that computation time increases with the size of interrogation window for the SSD method, which would be similar to extension of the interrogation window to an extra dimension. However, we anticipate that application of QPV to 3D QPI could be used to resolve transport in 3 dimensions and study thicker samples such as rounded cells and tissue slices. Labelling of different organelles in the cells using 3D QPI can reveal the distribution of organelles through cell height more accurately, enabling separation of cell compartments such as the nucleus from the cytoplasm, or quantification of transport in individual organelle systems, thus further revealing information about interaction of different organelles within cells.

QPV has a number of advantages over alternative approaches for studying mass transport within cells. Unlike many other label-free methods, QPV automatically tracks intracellular features based on PIV. Additionally, as a label-free method, QPV avoids issues with phototoxicity, photobleaching, or label dilution over time, while still giving results for intracellular diffusivity that are comparable in magnitude to methods requiring labels. Another major advantage of QPV is that it uses the same analysis for both nucleus and cytoplasm, allowing direct comparisons between these two compartments that are difficult to label consistently with other approaches. Overall, this work suggests that QPV is a valuable tool to study intracellular transport and biophysics.

## METHODS

### Microscopy

We performed QPI with an Olympus IX83 inverted microscope (Olympus Corporation, Japan) in brightfield with a 100X, 1.3 numerical aperture oil-immersion objective, and 1.2X magnifier to match the Nyquist criteria for diffraction limited imaging with a QWLSI wavefront sensing camera ^29,47^ (Phasics SID4-4MP (Phasics, France) camera). 120 ms exposure with red LED illumination (623 nm, DC2200, Thorlabs, USA) was used for QPI image acquisition. We used MATLAB (Mathworks, USA) for automated image acquisition. We connected the illumination sources, Retiga camera and stage with MicroManager open-source microscopy software ^48^. The Olympus IX83 microscope (Olympus corporation, Japan) and Phasics camera were connected directly through MATLAB. QPI images were captured every 1 minute and fluorescence images every 30 minutes at 30 imaging positions in every dataset. A flipping mirror arrangement (IX3-RSPCA, Olympus Corporation. USA) enabled alternate fluorescent and QPI. X-Cite 120LED light source (Excelitas technologies, USA) and Retiga R1 camera (Cairn research Ltd, UK) at 300 ms exposure were used for the fluorescence, with an Olympus U-FBNA filter cube for green mAG fluorophore and Semrock mCherry-B-000 filter cube (IDEX health & science, USA) for imaging the red mKO2 fluorophore. Uniform 37°C and 5% CO_2_ conditions were maintained using an Okolab stage-top incubator (Okolab, Italy) and custom-built objective heating collar, temperature controlled by Thorlabs temperature controller (Thorlabs, USA). For live cell imaging, cells were plated in Ibidi µ-high treated dishes at 30% confluence. Cells were incubated two hours after seeding, before moving to microscope for imaging, and imaging was initiated 20 minutes after transferring cells to microscope stage to ensure uniform temperature distribution in dish. Four sets of 30 live cells were imaged every for 8 hours for every imaging session.

### Cell culture

Cell culture procedures complied with the University of Utah BSL-2 guidelines. MCF7 (mammary epithelium) cell lines donated by Welm lab (HCI, Utah) were cultured in Dulbecco’s modified eagle’s medium (DMEM) (Gibco™ 11330057, Thermo Fisher Scientific, USA) with 10% fetal bovine serum (FBS) (Corning™ 35015CV, Fisher Scientific, USA). mKO2-hCdt1 and mAG-hGem tagged FUCCI expressing RPE-1 cells from Edgar lab (HCI, Utah), prepared by Yiqin Ma, were cultured in Gibco DMEM with 10% FBS and 5% Penicillin-Streptomycin. Penicillin-Streptomycin was removed while imaging the cells. Cells were split from 1 to 6 to 1 to 3 ratio at confluence less than 80% at a frequency based on their growth rate.

### Cell FUCCI tagging

RPE cells donated by the lab of Bruce Edgar (University of Utah) were received FUCCI tagged with mAG-hGem and mKO2-hCDt1. Cells express the mKO2 tag on Cdt1 in the nucleus during G1 phase (red nucleus), added mAG tag on Geminin in the nucleus during G1 to S phase transition (yellow nucleus due to combination of red mKO2 and green mAG) and the loss of mKO2-hCdt1 with the onset of S phase leading to green fluorescent labelled nucleus in S and G2 phase. The fluorescent image of the FUCCI tagged nuclei captured every 30 minutes was segmented using the k-mean algorithm (available as a built-in function in MATLAB) and overlapped with the QPI image. We bin the cells in S and G2 phase by considering three hours immediately after the nuclear label turns green and three hours before cell cytokinesis as the G2 phase of cell cycle.

### Mycoplasma testing

We DAPI (Fisher-NC9677247) stained fixed cells, grown 80 to 90 % confluent, after permeabilizing the nuclei using methanol at 4ºC, to test for mycoplasma contamination. The DAPI stained cells were imaged using a Retiga R1 camera at 500 ms exposure time and Olympus U-FUNA filter cube, illuminated by X-Cite 120LED. We checked the presence of DAPI stained puncta outside the nucleus as an indicator for mycoplasma contamination.

### Initial processing of QPI data

The Phasics SID4BIO camera for QPI uses a modified Hartmann mask diffraction grating to compute the phase gradient in two orthogonal directions which were converted to phase measurements using the Phasics Matlab SDK (Phasics, France). Phase shift was converted to dry mass using an assumed specific refractive increment of 0.18 µm^3^/pg for cell material ^22^. The phase shift of light (*ϕ*) expressed as an optical path length is directly related to the dry mass (*m*) at a given pixel through the specific refractive increment (α) ^49^ as:

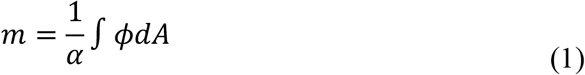

where *A* is the area of each pixel of the phase shift image. QPI images were then background corrected by fourth order polynomial curve fitting to regions in the phase image outside of cells.

### Fixed cell imaging

RPE and MCF7 cells were plated in Ibidi µ-high treated dish at 40% confluence. Cells were fixed by removing cell culture media, washing with PBS and incubating in 4% paraformaldehyde (PFA) at 37ºC for 10 minutes. The PFA was further removed, cells washed with PBS, refilled with fresh PBS and sealed and stored until imaging. During imaging, the dish was heated to 37ºC on Okolab stage-top incubator 20 minutes prior to imaging to avoid condensation. Cells were imaged at each 0.05 µm step stage translation in the vertical direction.

### Image registration

#### Sum of squared differences (SSD)

SSD was then performed on overlapping discretized interrogation windows of 15 by 15 pixels, spaced by 1 pixel, in the background corrected QPI image. The SSD of two image regions was computed as:

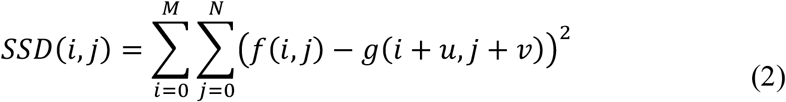

where *f* and *g* are the two interrogation windows from successive images to be matched using SSD with *i* and *j* the location of pixels comprising the image, *M* and *N* are the width and length of the image in pixels and *u* and *v* are the displacement introduced at each iteration of the SSD calculation.

On expanding the SSD equation (2) we obtain:

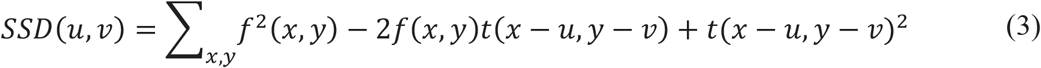

The second term in equation (3) contains the cross-correlation while the third term is related to the nonuniform energy density of QPI data. SSD, therefore, can be thought of as a method of normalization of the cross-correlation.

#### Normalized cross correlation (NCC)

NCC between interrogation windows was measured using the MATLAB function *normxcorr2*. The function computes the cross correlation in the Fourier plane to improve computation efficiency based on the equation of normalized cross correlation.

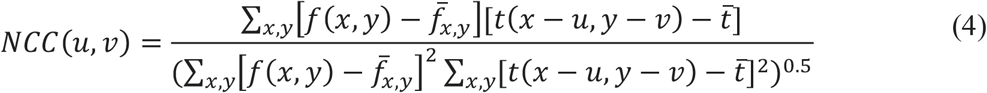

Here, f and t are the images and 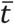 and 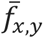 are the mean of the images. The displacement of the highest pixel in the normalized correlation plane from the center pixel was used to compute the displacement of each interrogation window. NCC, similar to SSD, normalizes the cross-correlation plane by division with image auto-correlation.

#### Optical flow reconstruction (OFR)

OFR developed by Lucas and Kanade ^50^ measures the transport of intensity profile in images, given by

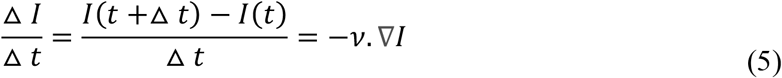

Here *I(t+Δt)* and *I(t)* are intensity of images time *Δt* apart, in our case 1 minute. We used the code available from ^32^ for OFR displacement calculation. The *BlurSize* parameter for blurring image using Gaussian blurring before OFR in code was set to 1. The *BlurStd* parameter was set to 1,2,5,8 and 12 for understanding impact of Gaussian blurring (figure S4 c and d).

#### Mutual Information (MI)

MI was computed using a method implemented in MATLAB that extracts mutual information from the joint histogram and joint entropy of the registered images that was adapted to perform mutual information (MI) image registration calculation on sliding interrogation windows inside grayscale QPI images ^51^.

### Quantitative phase velocimetry

Our implementation of QPV is based on the SSD image registration method ^34^ applied to 15 by 15 pixel (3.57 by 3.57 µm) interrogation windows within cells. A magnified view of one 15 by 15 pixels interrogation window inside an example cell difference image (**figure 1c**) shows the movement of individual cell components (pointed by red arrow) to a position on the left side of the window, pointed by black arrow (15 by 15 pixel inset in **figure 1c)**. SSD produces the lowest sum of squared difference when such patterns of mass within an interrogation window overlap between two displaced windows from successive imaging frames (**figure S1a)**. Gaussian fitting on the 3 by 3 pixel region neighborhood around the lowest value of the computed SSD gives sub-pixel localization of displacements (**figure S1b)**. This results in the measured displacements that form the basis of QPV (**figure S1c**). Spurious velocities were removed using a conditional median filter, thus retaining the original values of subcellular velocity computed. From the known time gap between frames set during cell imaging, we compute the velocity of mass transport from these displacement measurements (**figure 1d**). Processing was performed using computational resources allocated by the Center for High Performance Computing (CHPC) at the University of Utah. Code is available on GitHub (https://github.com/Zangle-Lab). The relative computational time of each image registration method (**figure S4**) was estimated by computing intracellular velocity maps of 10 RPE fixed cell images at each set of interrogation window sizes on a general-purpose computer with a 6 CPU core Intel i7 8700K, 3.7GHz processor and 16 GB RAM.

### Calculation of measurement error and velocity direction

The error of displacement magnitude (*E*_*mag*_) was calculated as:

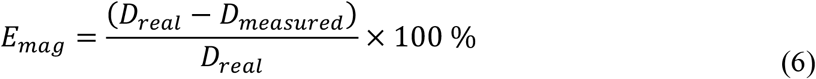

where *D*_*real*_ is the actual magnitude of displacement introduced by movement of stage, and *D*_*measured*_ is the average measured displacement magnitude. *D*_*real*_, velocity magnitude, is measured from x component (*V*_*x*_) and y component (*V*_*y*_) of velocity as:

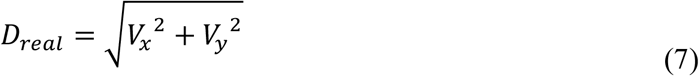

Direction error (*E*_*dir*_) was computed as the angle between the computed and actual displacement vector:

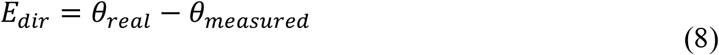

*θ*_*real*_ is the actual angle of displacement introduced by stage and *θ*_*measured*_ is the average angle measured by image registration method. Is calculated from x component (V_x_) and y components (V_y_) of velocity using four quadrant inverse tangent:

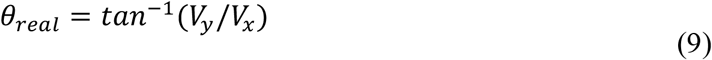

### QPV error model

Power spectrum (*Pf*) of image (*Image*) was calculated by

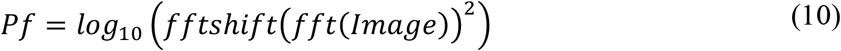

*fft* is the Fourier transform of the image and *fftshift* rearranges the Fourier transform to shift the zero-frequency component to the center of the array. Power spectra of the cell images was azimuthally averaged to convert power density image to 2-dimensional plot of power density vs frequency.

We estimated the optical diffraction limit as:

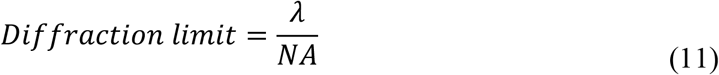

where *λ* is the wavelength of light and *NA* is the numerical aperture of the objective used for imaging, assuming that the illumination NA = 0 to generate a spatially coherent plane wave for QPI^29^. The code for constructing the particle size dependent error model is also available on GitHub (https://github.com/Zangle-Lab).

We perform QPV intracellular velocity measurement on 2D QPI images. Thus, the features outside the depth of the field of the microscope will be out-of-focus, and thus blurred. The depth of field (*DOF*) for illumination with incoherent light can be calculated by^52^,

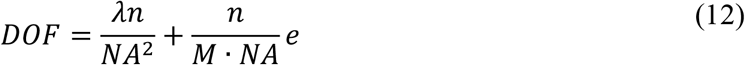

Where *λ* is the wavelength of light, *n* is the refractive index of media between the objective and the sample, *NA* is the numerical aperture of the objective, *M* is the magnification, and *e* is the camera resolution at the CCD plane, here used as the size of a phase pixel. Equation 5 gives a depth of field of 0.85 µm. However, we use near-coherent illumination by closing the condenser as far as possible (NA_condenser_ ≈ 0). Therefore, the depth of field of our system is approximately double the depth of field calculated by equation (12)^52^, *i*.*e*., 1.7 µm.

OPL from is computed as,

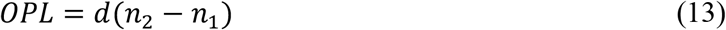

Where *d* is the actual distance, *n*_*2*_ is the refractive index of the sample and *n*_*1*_ is the refractive index of the surrounding media. The refractive index of DMEM cell culture media (*n*_*1*_) is 1.337^53^ and general refractive index used for cells (*n*_*2*_) is 1.36^54^. The interrogation window power density is calculated from the power spectrum distribution in the interrogation windows inside cells. Interrogation window power density is computed as the area under the 2D power density curve as an indication of feature density in QPI data.

Synthetic data was generated by plotting circles of specified radius at random locations. Poisson noise of mean 5.5, scaled down by 1e12, was added to the synthetic data images. The background of the images was fit using polynomial fitting and then flattened by subtracting this from the synthetic data, the same procedure as applied to real QPI data. The background of the images that remains thus resembles Perlin noise. This was confirmed by the similarity of pattern of the power spectrum image of the created background to the expected Perlin noise power spectrum image. For select images (figure S8), original QPI background noise was added to synthetic data to assess its utility in matching the power spectra of QPI data. This was performed by addition of empty background corrected QPI images to the synthetic data.

### Deformation velocity

The whole cell velocity was first computed by tracking the centroid of the entire segmented cell outline. The deformation velocity, a measure of the magnitude of local deformation, was then computed as the magnitude of the difference between the control volume and cell centroid displacement vectors. To determine nuclear vs. cytoplasmic deformations, the cell nucleus was segmented from cytoplasm using k-means segmentation algorithm on FUCCI fluorescence images. Cytoplasm and nucleus mask were used to separate subcellular 3-minute displacement tracks in corresponding regions and used to compute cytoplasm and nuclear velocities separately.

### MSD analysis

Relation between MSD and time lag (*τ*) is given by

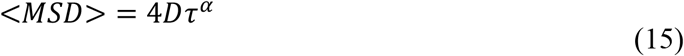

D is the diffusion coefficient and *α* is the anomalous constant. The proportionality constant is 4D as the tracking is two-dimensional. The effective particle size (*d*_*p*_) is obtained from power spectrum of QPI image of the cell, weighting the particle size by the power spectrum density, within the limit of particle sizes visible by the interrogation window size.

### Statistics

Welch’s two-tailed t test with unequal variances was used to calculate the significance of differences between experimental groups. Error bars are reported as the standard error of the mean based on the number of cells in each test group.

## ACKNOWLEDGMENTS

This work was supported by the University of Utah and NIH K25CA157940. We thank Dr. Bruce Edgar’s lab, Dr. Yiqin Ma at the Huntsman Cancer Institute, University of Utah for donation of FUCCI tagged RPE cells. We also thank Dr. Alana Welm’s lab at Huntsman Cancer Institute, University of Utah for donation of MCF7 cells. Finally, the support and resources from the Center for High Performance Computing at the University of Utah are gratefully acknowledged.

## AUTHOR CONTRIBUTIONS

Conceptualization: TZ; Experiments: SP; Theoretical formulation: SP, TZ; Funding acquisition: TZ; Investigation: SP, TZ; Code: SP, TZ; Supervision: TZ

## ADDITIONAL INFORMATION

The authors declare that they have no competing interests.

