## Supplemental Material for "Quantitative phase velocimetry measures bulk intracellular transport of cell mass during the cell cycle"

**The PDF file includes:**

Supplementary figures 1 to 20
Legends for movies S1 and S2

**Other Supplementary Material for this manuscript includes the following:**

Movies S1 and S2

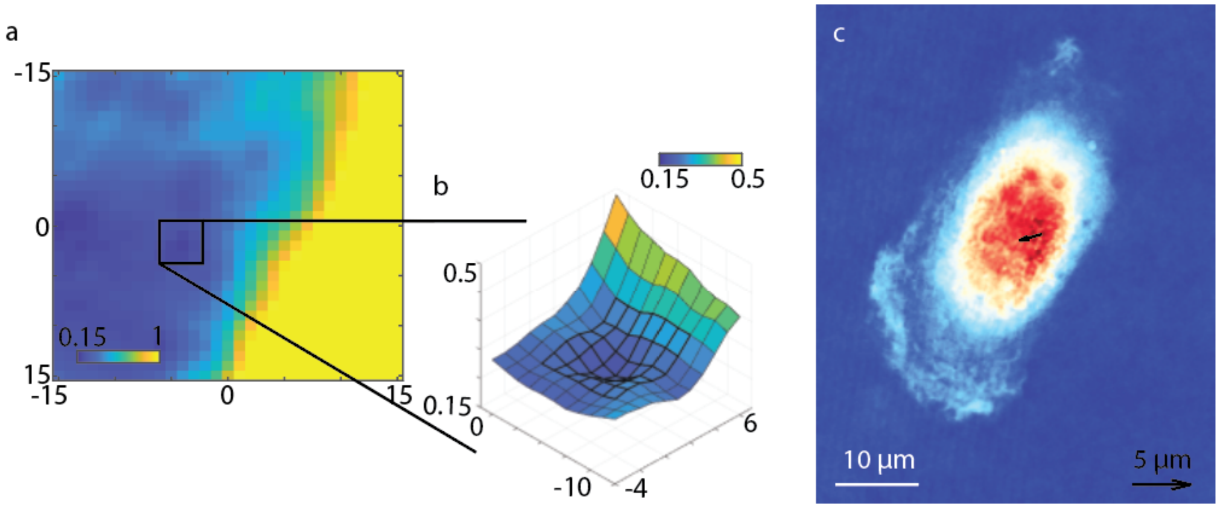


**Figure S1: SSD for one interrogation window in QPI image. (**a) QPV uses sum of squared difference (SSD) image registration method, which computes the sum of square of the difference image of interrogation windows pixels at different displacements. (b) Local Gaussian fitted surface in 3 by 3 neighborhood of the lowest pixel on SSD. (c) Dry mass distribution of the cell with displacement inside the interrogation window indicated as a black arrow (5 µm velocity scalebar for comparison).

a

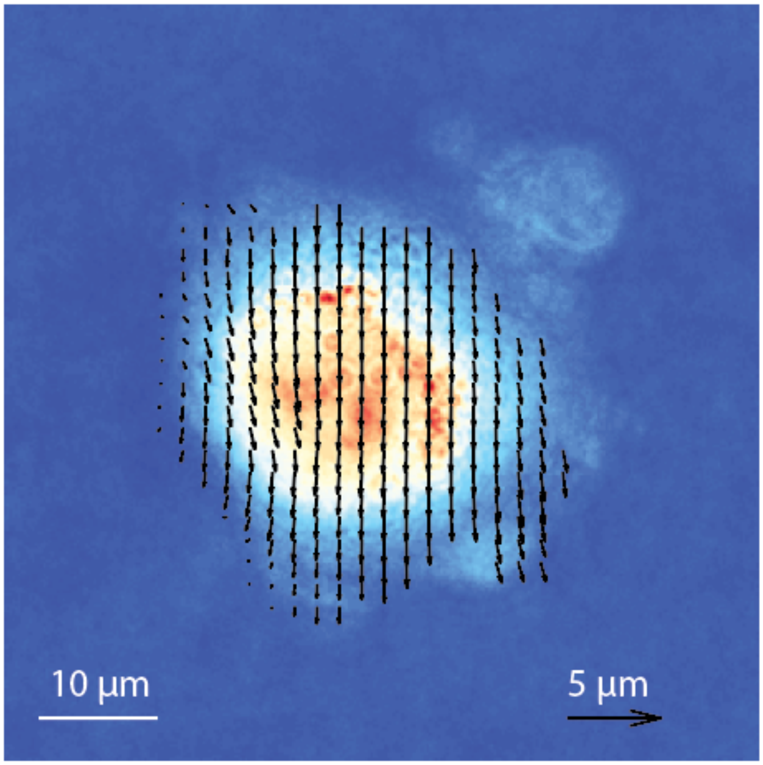


**Figure S2: QPV validation uses artificially moved fixed cells** (a) QPV on fixed MCF7 cell, moved 1.5 µm downwards artificially, measures the intracellular displacement.

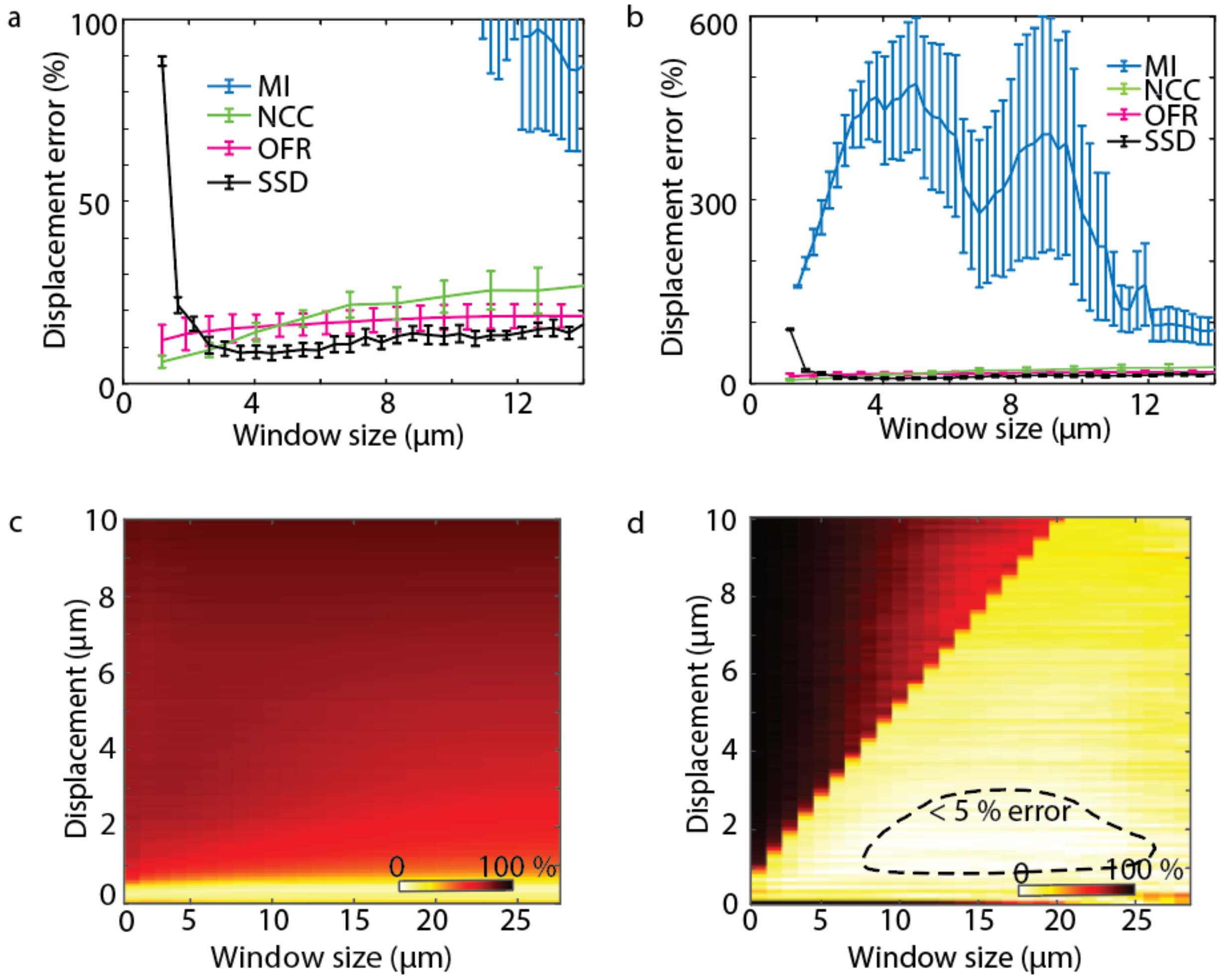


**Figure S3: Performance of SSD, OFR, MI and NCC on MCF7 cells.** (a) Comparison of mutual information (MI), normalized cross correlation (NCC), optical flow reconstruction (OFR) and sum of squared difference (SSD) for intracellular velocity computation on MCF7 QPI data at different interrogation window size shows SSD has highest accuracy at all window sizes ($n$ = 9, error bars show standard error of the mean). (b) Full range of data in (a) shows velocity error using MI with small window sizes. (c) Velocity accuracy measurements as a function interrogation window size and displacement of MCF7 fixed cell using OFR shows very narrow range of displacement measurements possible by OFR with acceptable accuracy ($n$ = 9). (d) QPV velocity measurement accuracy as a function interrogation window size and displacement shows less than 10% error with an interrogation window size less than the displacement to be measured and a region of < 5% error (dashed line, $n$ = 9).

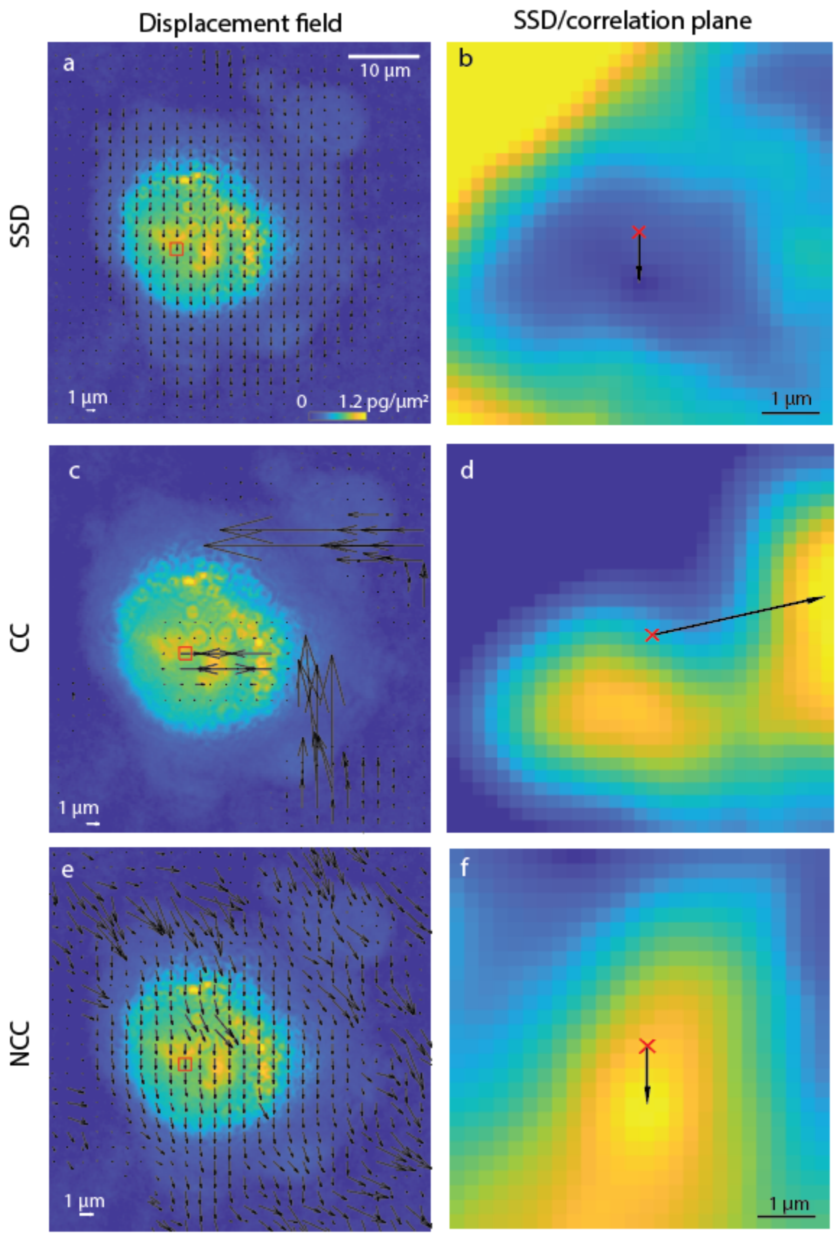


**Figure S4: Intracellular velocity estimation using SSD, CC and NCC shows SSD is similar to NCC.** (a) Displacement measured using SSD in an artificially moved fixed cell shows uniform displacement distribution of magnitude 1 µm (4.2 pixels). (b) SSD plane from one 15 by 15 pixel interrogation window (marked by red square in panel a) inside the cell in (a) shows the minimum value is 4 pixels away from the center of the plane (marked with a red x). (c) Intracellular velocity in the fixed cell computed using cross-correlation shows significant velocity errors. (d) Cross-correlation plane computed for same interrogation window as in (b) shows erroneous 16 pixel displacement of maximum value from the center of the cross-correlation plane. (e) Intracellular velocity estimated using normalized cross-correlation. (f) Correlation plane from interrogation window in the red box shows 4 pixel displacement from the center pixel.

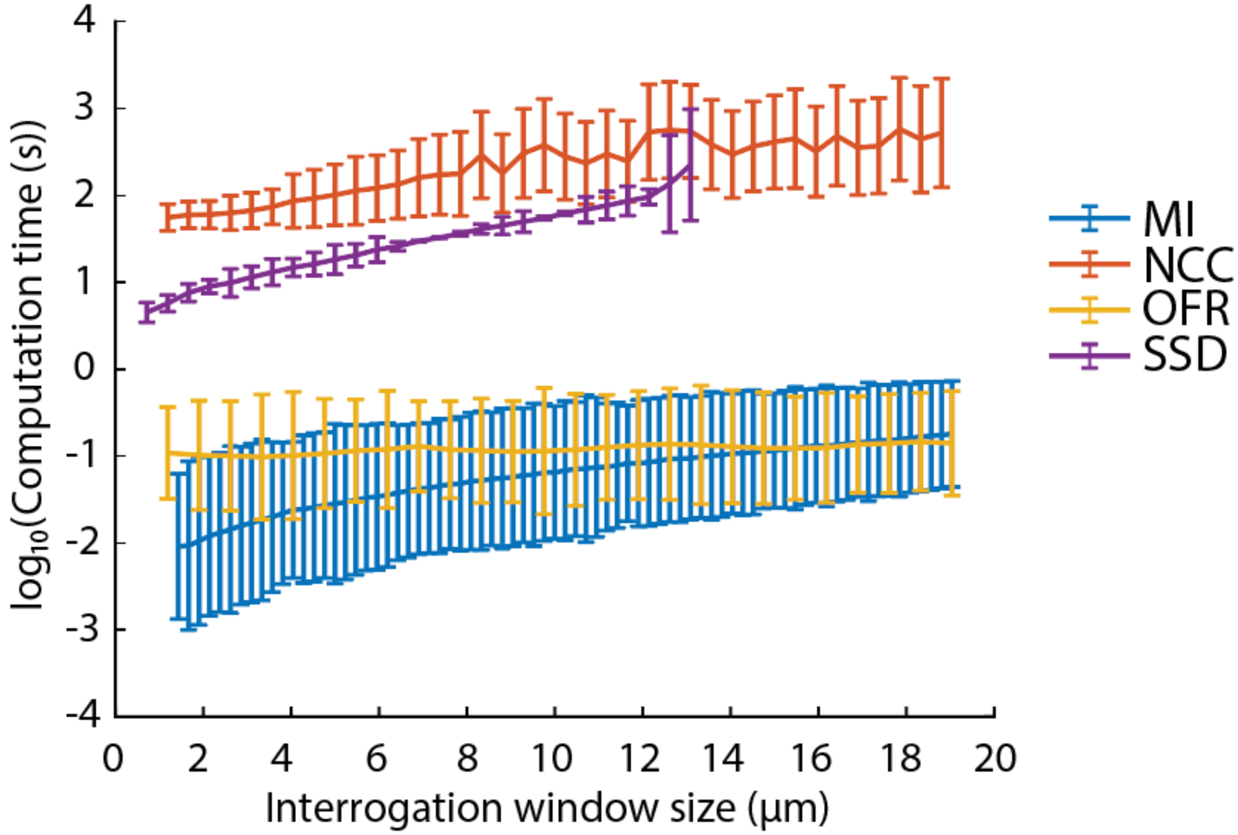


**Figure S5: Computation time of MI, OFR, SSD and NCC.** Semi-logarithmic plot of computation time of intracellular velocity map at different window sizes for MI, OFR, SSD and NCC for 10 RPE fixed cell images. Computation time for NCC and SSD is larger than OFR and MI. Intracellular velocity computation time for SSD increases with increasing window size, showing most computational efficiency at smallest window sizes.

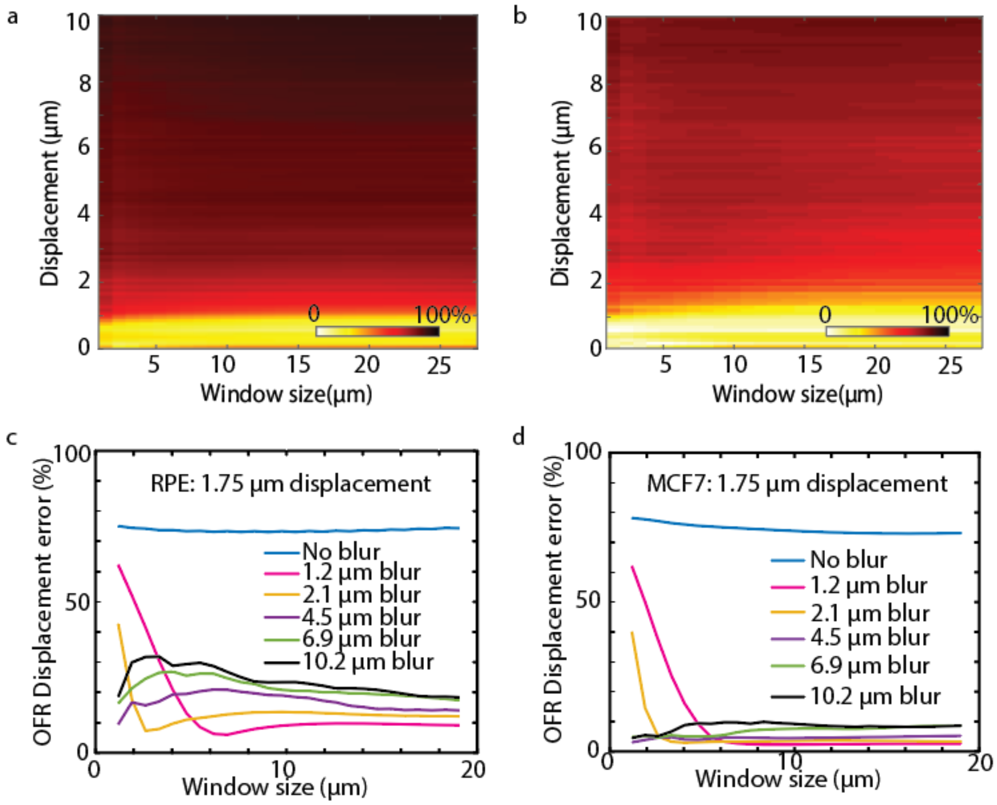


**Figure S6: The performance of OFR is moderately improved by Gaussian blurring of QPI images.** (a) OFR displacement estimation error at different window sizes without Gaussian blurring. (b) OFR displacement error at different window sizes with 1.2 µm standard deviation Gaussian blur. (c) OFR displacement error for RPE cells displaced 1.75 µm for Gaussian blur from 0 to 10.2 µm. (d) OFR displacement error for MCF7 cells displaced 1.75 µm for Gaussian blur from 0 to 10.2 µm.

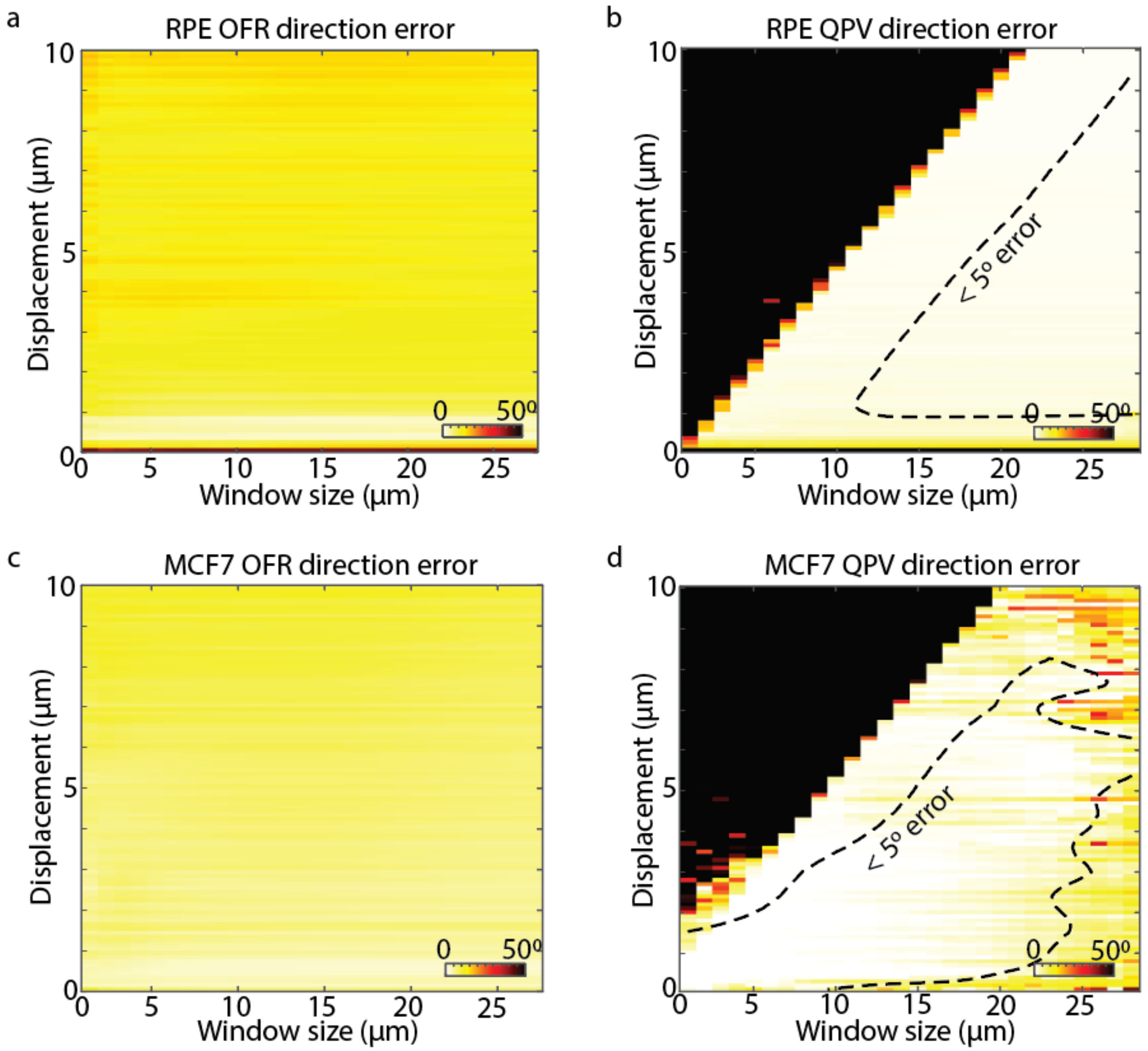


**Fig S7: Comparison of OFR and QPV velocity direction error.** Averaged absolute value of (a) OFR and (b) SSD direction error as a function of displacement and window sizes in RPE cells. Averaged absolute value of (c) OFR and (d) SSD direction error as a function of displacement and window sizes in MCF7 cells. RPE $n$ = 11, MCF7 $n$ = 9.

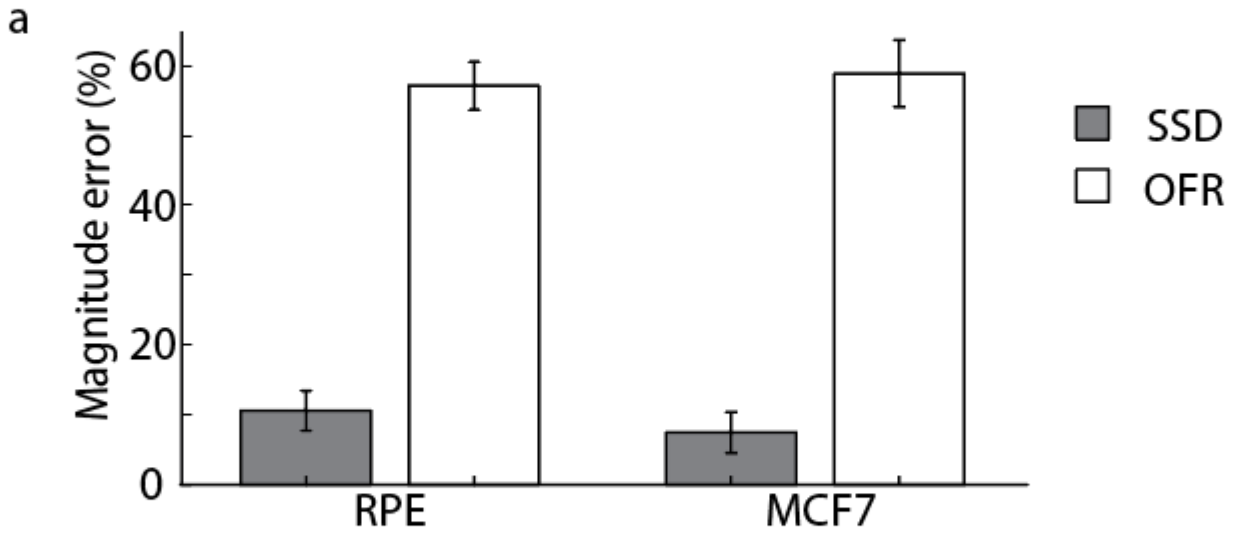


**Figure S8: SSD has lower error than OFR at the operating window size.** (a) Displacement magnitude error for 15 x 15 pixel interrogation window size averaged over $n$ = 11 RPE and $n$ = 9 MCF7 cells, computed with SSD and OFR methods. Error bars show standard error of the mean.

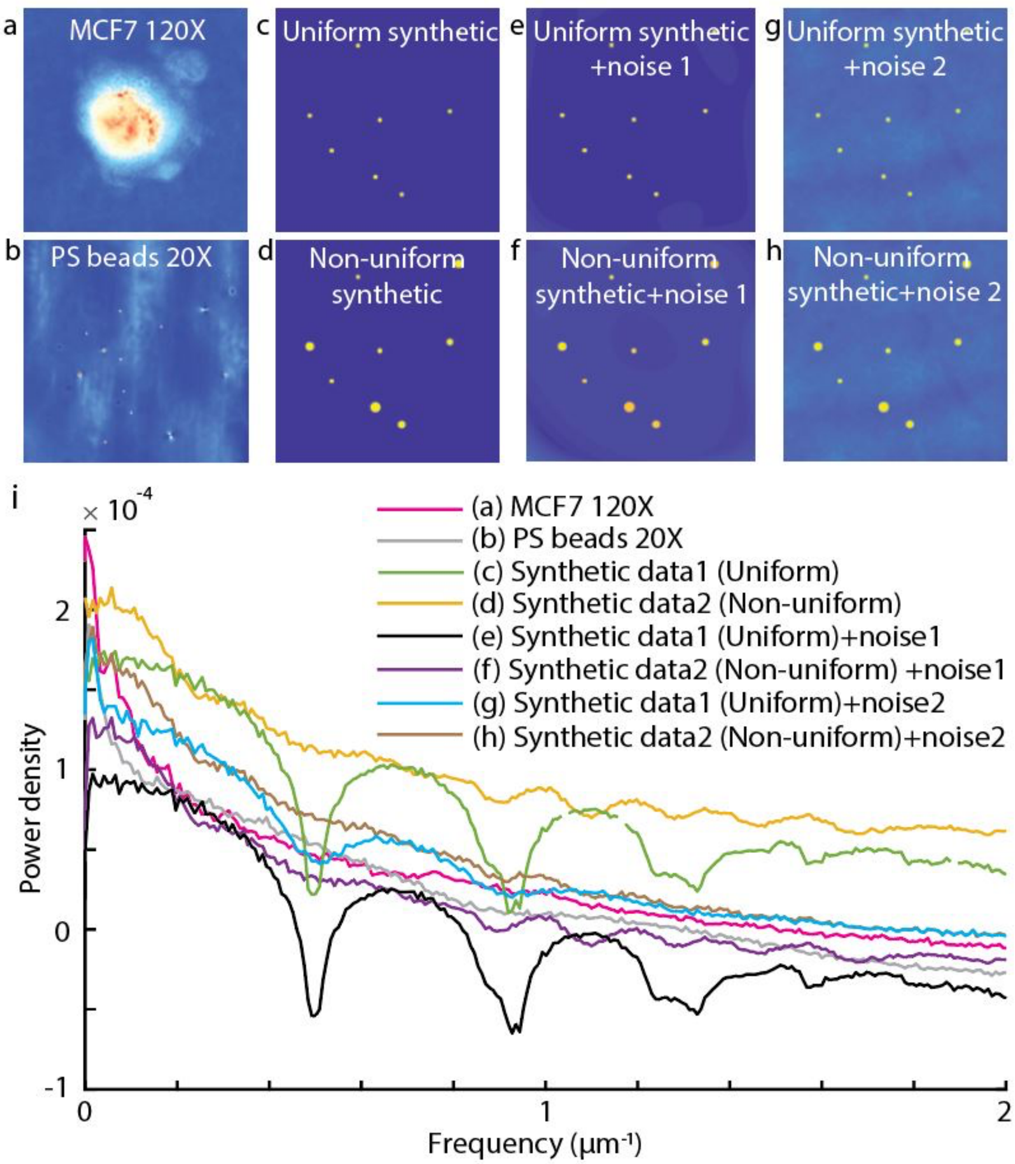


**Figure S9: QPI data characterization comparing to synthetic data** (a) Comparison of background corrected QPI image of (a) MCF7 fixed cell at 120 X. (b) Polystyrene beads at 20X. Synthetically generated (c) uniform and (d) non-uniform circle data, with added Perlin noise (e and f). (g) Synthetically generated uniform data with QPI image background. (h)Synthetically generated non-uniform circle data with the QPI image background. (i) Power spectrum of the MCF7 QPI data at 120x (magenta), 20x polystyrene bead QPI data (gray), uniform synthetic data without (green) and with noise (black), non-uniform synthetic data without (yellow) and with (purple) noise, synthetic uniform data with QPI background noise (blue) and synthetic non-uniform data with QPI background noise (brown), shows similarity of QPI data to non-uniform synthetic data. The power spectrum of the uniform-sized circle data shows narrow ranges of spatial frequencies characterized by the size of the beads and the space between them while the non-uniform circles have a smoother power spectrum reflecting the presence of a wider range of spatial frequencies

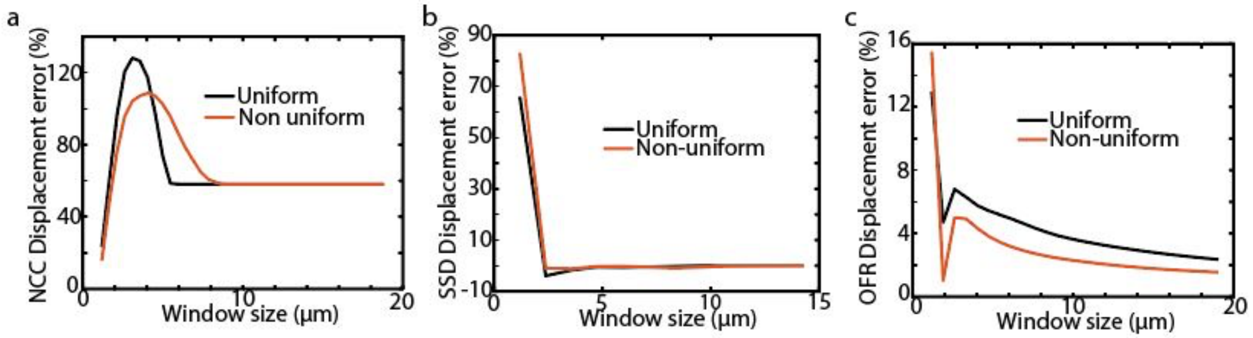


**Figure S10: NCC, OFR, and SSD performance for synthetic data with different characteristics at 0.1 µm displacement.** (a) NCC performance versus window size for uniform synthetic data (black) and non-uniform data (red). (b) SSD performance versus window size for uniform (black) and non-uniform synthetic data (red). (c) OFR performance versus window size for uniform synthetic data (black) and non-uniform synthetic data (red).

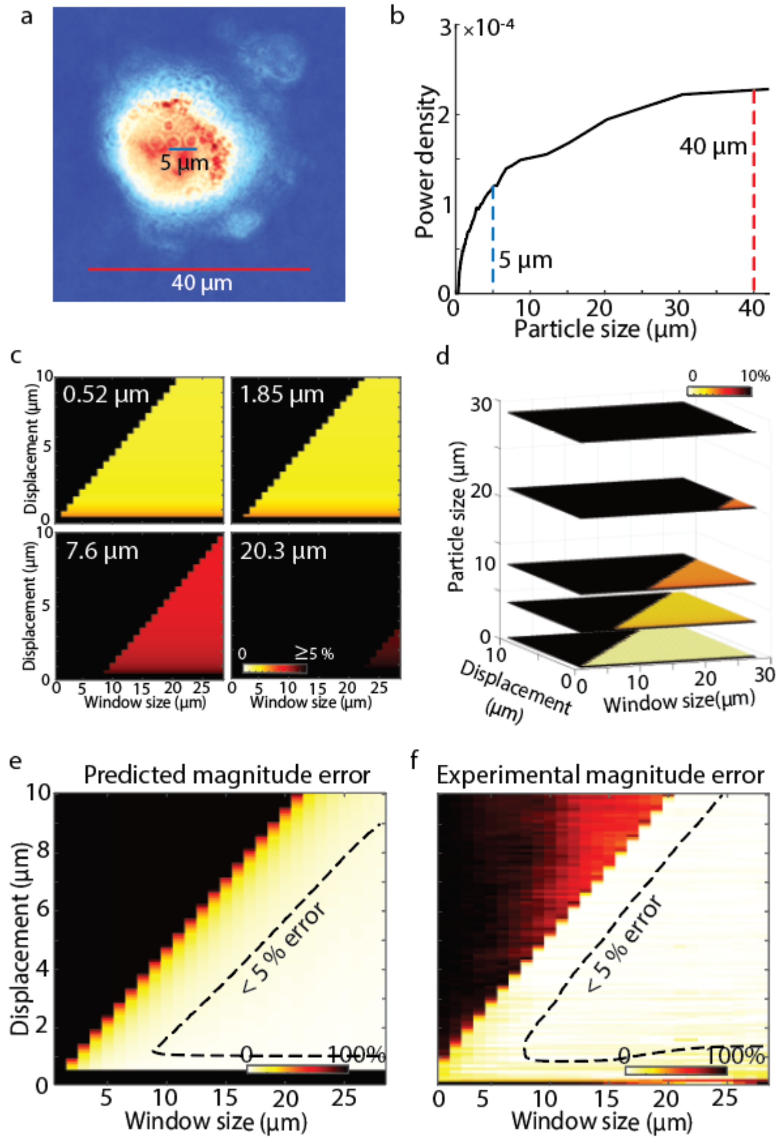


**Figure S11: Modelled velocity measurement error for QPV versus measured error for fixed MCF7 cells**. (a) 5 µm organelle structure inside 40 µm MCF7 fixed cell. (b) Power density distribution of particle sizes of the MCF7 fixed cell with indication of power density of 5 µm and 40 µm structures. (c) Theoretical velocity estimation error at four particle sizes as a function of displacement and window size. (d) Velocity magnitude estimation error as in (c) shown as 3D projection with slices at 0.5, 5, 10, 20 and 30 µm particle size. (e) Averaging the data from (c-d) weighted by particle sizes from (b) gives an estimated error which is in agreement with the (f) measured velocity magnitude error of the same MCF7 fixed cell.

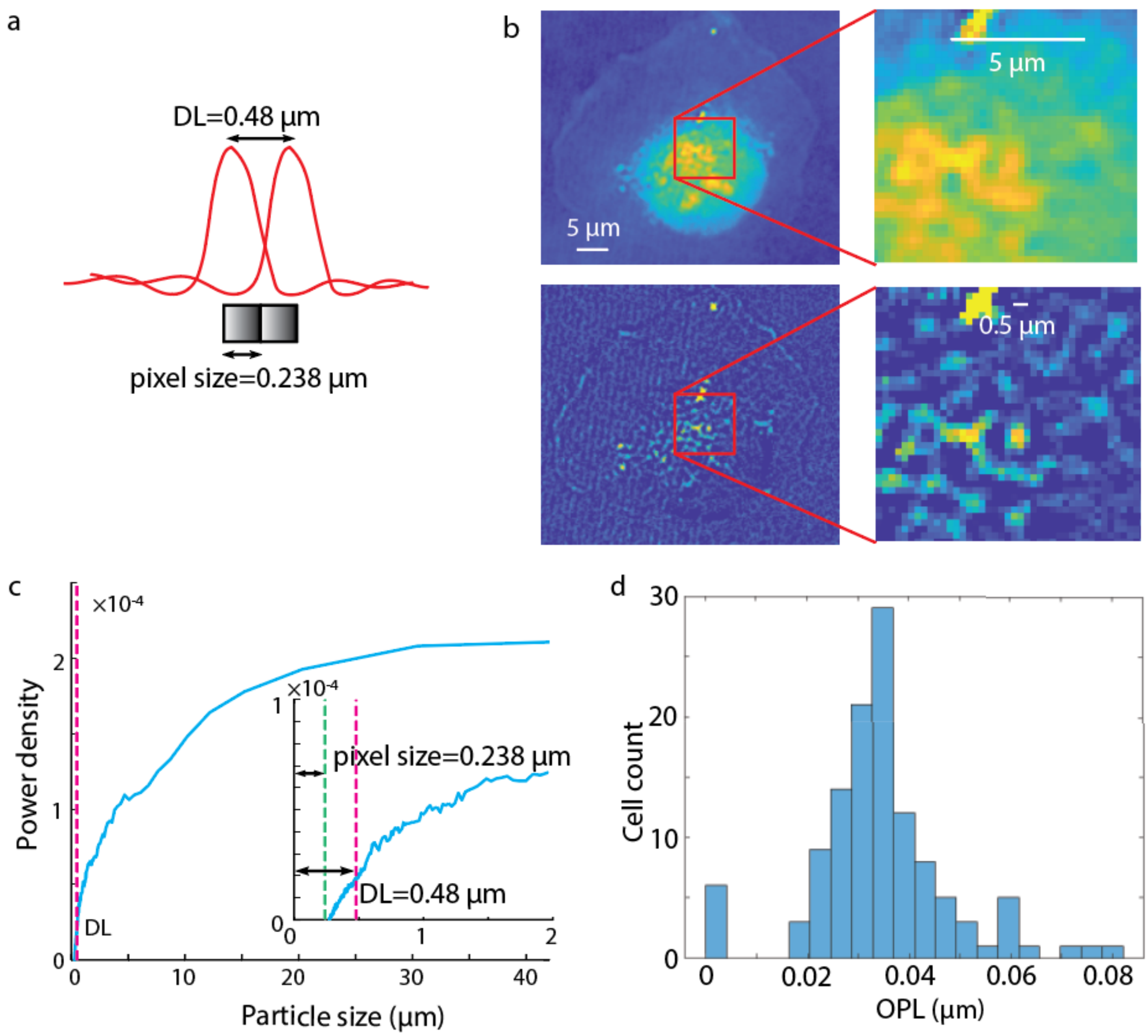


**Figure S12: QPI optical system satisfies the Nyquist criteria giving diffraction limited image resolution for QPV**. (a) The size of computed phase pixels is half of the diffraction limit of light in our system, computed as $\lambda/(NA_{illum} + NA_{objective})$, where $\lambda = 623$ nm, $NA_{objective} = 1.3$, and $NA_{illum} \approx 0$ because the condenser aperture is closed to approximate illumination with a plane wave. (b) shows both the raw QPI image of a fixed RPE cell (top) and a rolling ball filtered image (kernel diameter = 2 pixels, 0.48 µm) demonstrating the appearance of particle sizes at approximately the diffraction limit of the system visible in our QPI data. (c) The power spectral density of the cell from (b) shows the diffraction limit relative to the range of particle sizes used as features for QPV. (d) Histogram of maximum optical path lengths (OPLs) in RPE cells.

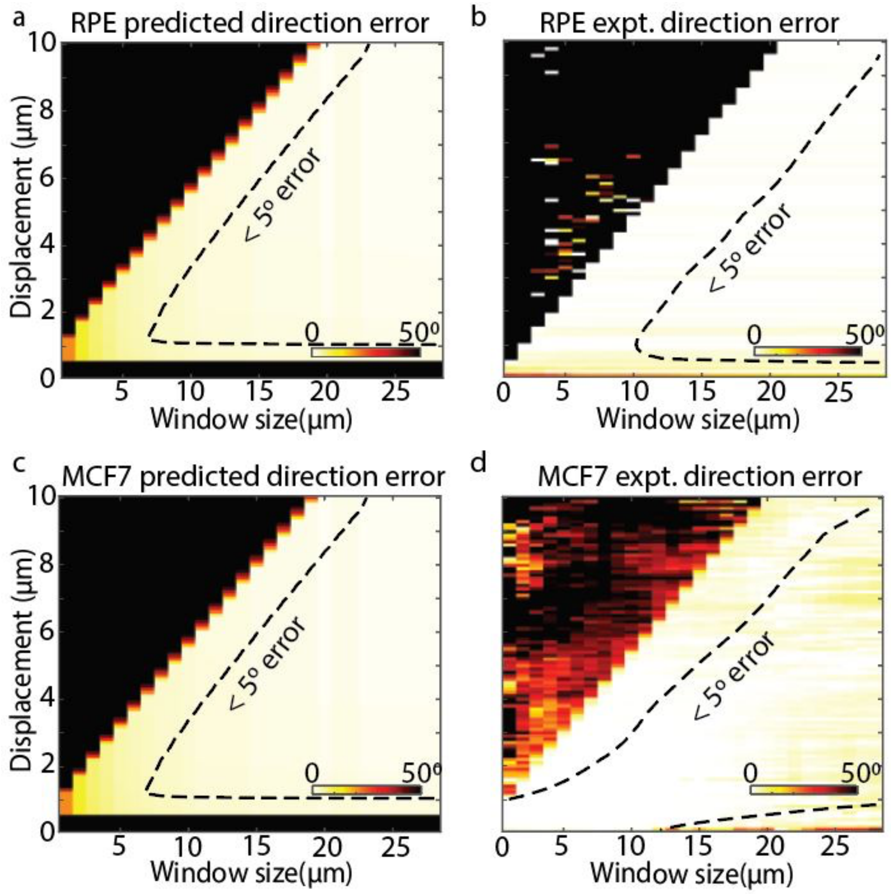


**Figure S13: Predicted QPV direction error agrees with measured QPV direction error for individual RPE and MCF7 cells.** (a) Predicted and (b) measured QPV direction error as a function of displacement and window size for an individual fixed RPE cell imaged at 120x. (c) Predicted and (d) measured QPV direction error as a function of displacement and window size for an individual fixed MCF7 cell imaged at 120x.

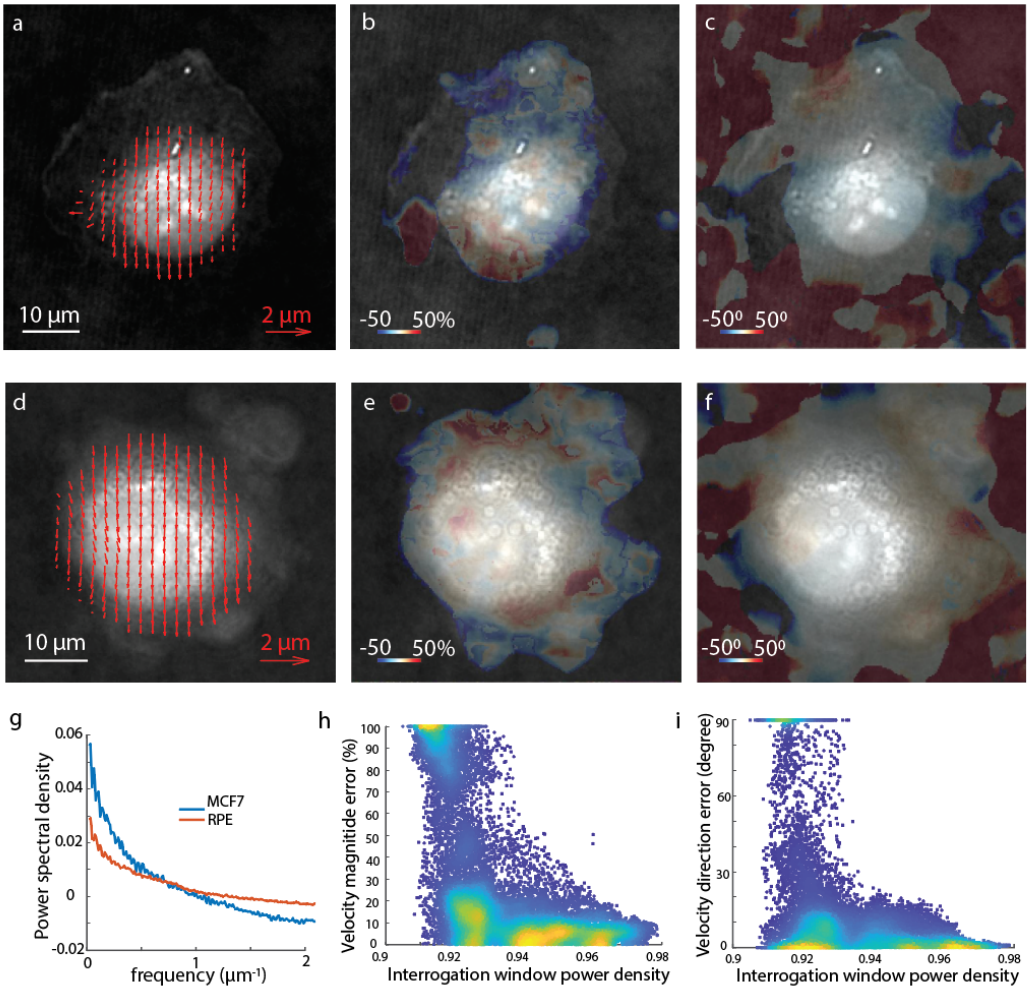


**Figure S14: Velocity estimate magnitude and direction error in fixed RPE and MCF7 cell shows higher error in regions with lower feature density.** (a) Intracellular velocity estimated using QPV in fixed RPE cell displaced 0.5 µm. Scalebar shows 10 µm. (b) Intracellular velocity magnitude error distribution on the fixed cell (a). Colorbar shows velocity magnitude error in percentage. (c) Deviation in estimated intracellular velocity direction from expected direction. Colorbar indicates the direction error in degrees. (d) Intracellular velocity in MCF7 fixed cell estimated by QPV. (e) Error in velocity magnitude in the MCF7 fixed cell shown in (d). Colorbar shows velocity magnitude error in percentage. (f) Direction error of the intracellular velocity distribution. Colorbar indicates direction error in degrees. (g) 2D power spectral density vs frequency plot of MCF7 cells (blue) and RPE cells (red). (h) Velocity magnitude error and (i) direction error vs particle power density in interrogation windows (a measure of feature density) shows a trend indicating lower error with increased particle power density. Interrogation window power density was computed as the area under the curve of the exponential of power spectra of individual windows.

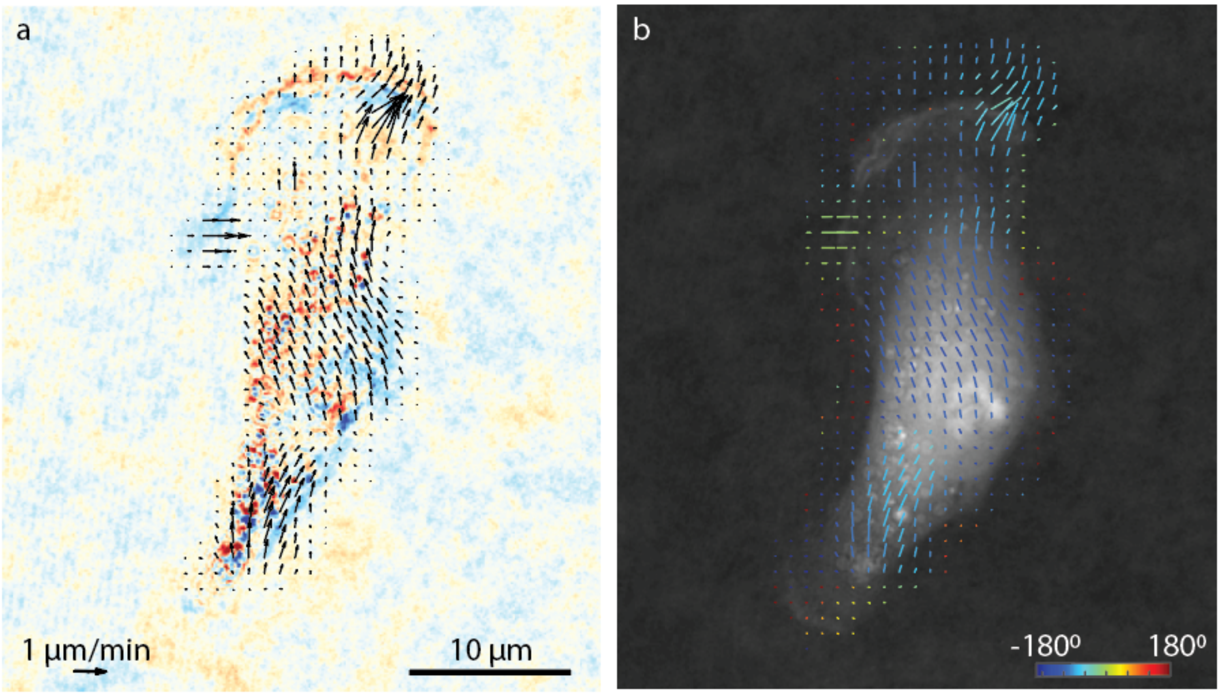


**Figure S15: QPV measures the magnitude and direction of long-range cell edge movements as well as short ranged movements.** (a) Difference between images at 1 min interval shows faster cell edge movement at top edge of cell, moving upward with cell center movement from left to right. (b) QPV also quantifies the direction of the movement in all areas of cell. Color of vectors indicates the direction of velocity in degrees, with 0° as a vector pointing to the right in these images.

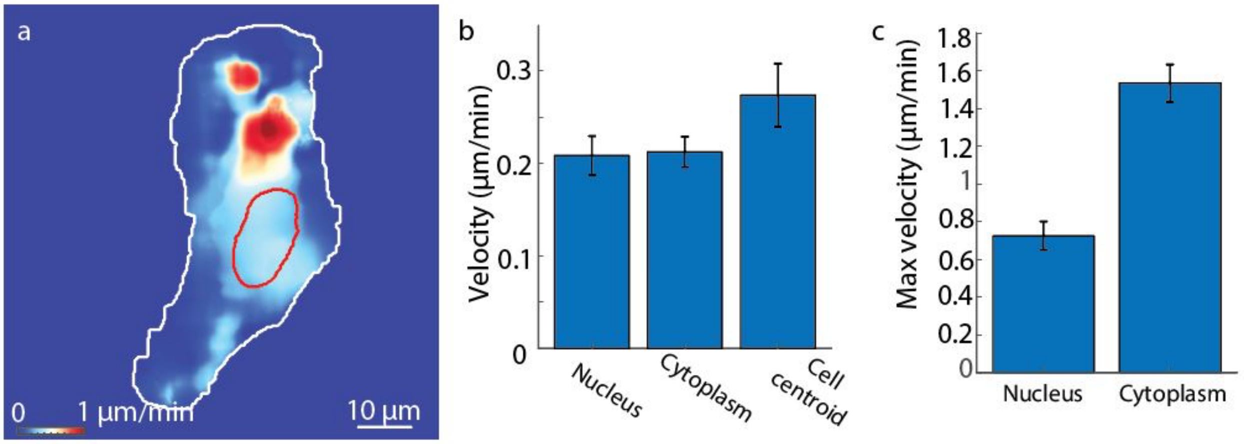


**Figure S16: QPV measures the average nucleus and cytoplasm velocity in live RPE cells.** (a) Average absolute intracellular velocity in a live RPE cell over 30 minutes. (b) Average velocity in nucleus and cytoplasm of RPE live cells is similar to the velocity of the whole cells measured using cell centroid tracking. Error bars show standard error of the mean, $n$ = 59.

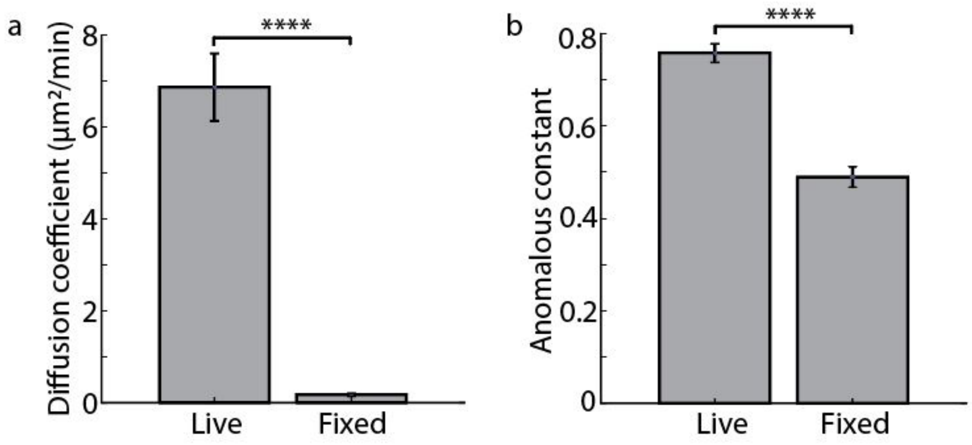


**Figure S17: Comparison of MSD analysis in live and fixed RPE cells.** (a) Effective diffusion coefficient from MSD analysis of live ($n$ = 119) and fixed ($n$ = 32) cells shows 40-fold smaller diffusion coefficient in fixed cells as expected. (b) Anomalous constant in fixed cell is moderately reduced relative to live cells. Error bars show standard error of the mean, ** $p < 0.01$, **** $p < 1x10^{-4}$

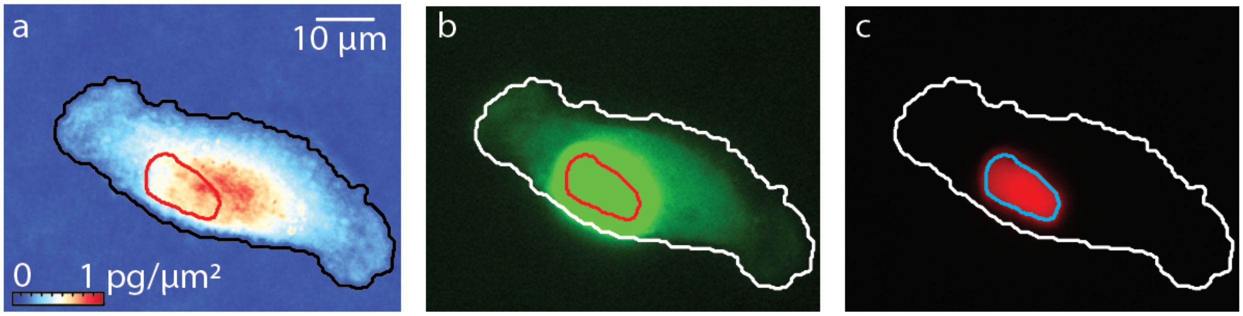


**Figure S18: Alignment of cytoplasmic and nuclear segmentation with FUCCI marker and QPI on individual RPE cell in S phase**. (a) Nuclear (red) and cytoplasmic (black) boundary on an RPE QPI image. Colormap shows dry mass density and scalebar indicates 10 µm. (b) The nuclear and cytoplasmic boundary superimposed on mAG fluorescence from FUCCI marker. The upper limit of the colormap has been scaled to the level of background fluorescence to show the alignment of cytoplasm boundary with the stray green fluorescence in cytoplasm of cell. (c) Nuclear and cytoplasm boundary on the mKO2 FUCCI marker fluorescence image showing alignment of nuclear segmentation.

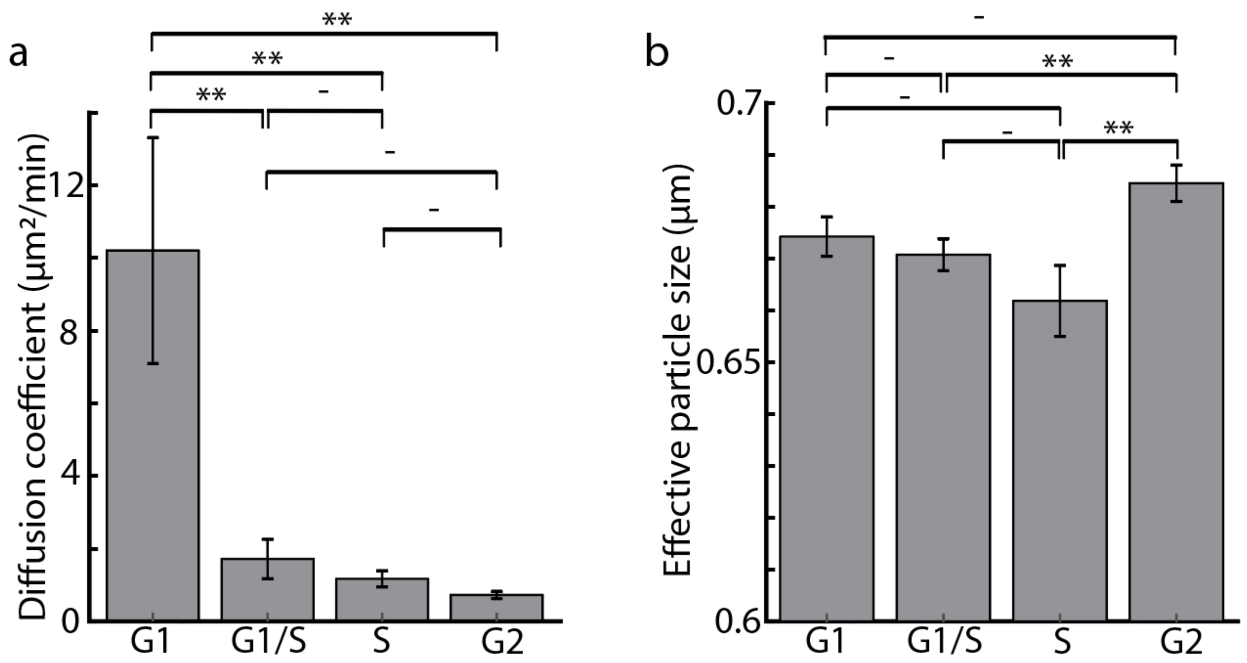


**Figure S19: Effective diffusion coefficient and particle size in live RPE cells.** (a) Whole cell average diffusion coefficient in RPE cells versus cell cycle phase. (b) Effective particle size calculated from power spectrum of populations of RPE cells in G1, S and G2 phases of cell cycle. Error bars show standard error of the mean, $n$ = 119 cells, $^{**}p < 0.01$.

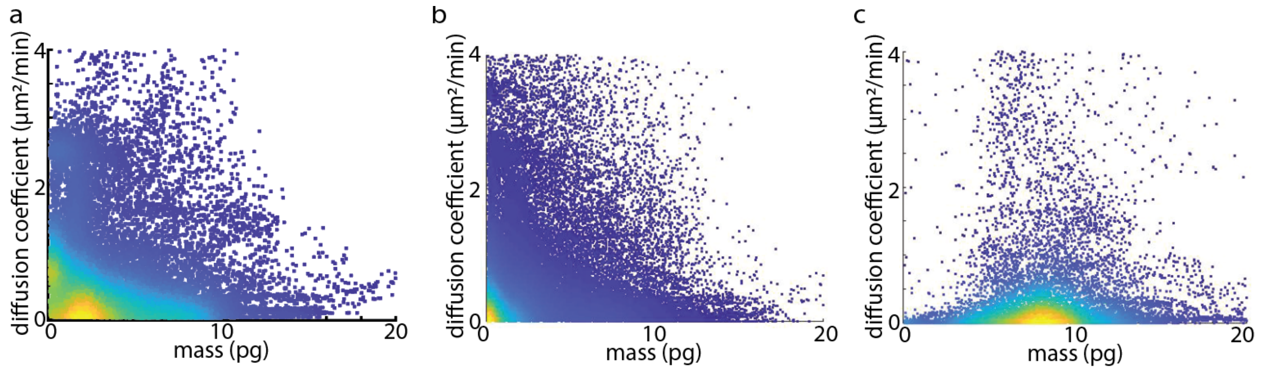


**Figure S20: Diffusion mass relation of control volumes in nucleus and cytoplasm of RPE cells.** (a) Diffusion coefficient versus mass for all regions tracked inside RPE cells. (b) Diffusion coefficient to mass relation in cytoplasmic volumes of RPE cells. (c) Diffusion coefficient vs mass in nuclear volumes in RPE cells.

**Movie S1.**
Deformation of grid overlayed on the RPE cell shown in figure 4 tracked with QPV for 30 minutes.

**Movie S2.**
Intracellular velocity map of RPE cell tracked with QPV over 30 minutes.
